# Cereal grafting in rice and pearl millet preserves photosynthetic performance and stomatal dynamics, establishing a platform for root-shoot communication studies

**DOI:** 10.64898/2026.06.25.734459

**Authors:** Crispus M. Mbaluto, Xabier Simón Martínez-Goñi, Anoop Tripathi, Pallavi Singh

## Abstract

- Cereal grafting using embryonic tissues has recently become technically feasible; however, the physiological consequences of cereal grafting remain uncharacterized.
- We systematically evaluate photosynthetic performance and stomatal dynamics across different graft combinations in two photosynthetically distinct species, rice (C_3_) and pearl millet (C_4_). We first assessed steady-state photosynthetic performance and dynamic stomatal responses in five-week-old rice and pearl millet grafts grown under a saturated water regime, to establish whether cereal grafting alters physiology at early stages. Next, we assessed same traits at the onset of optimal water regime, and after five days to determine whether any graft-induced effects on photosynthesis or growth persisted over time.
- We observed that across contrasting water regimes and at different plant developmental stages, cereal grafting did not alter growth, photosynthesis or stomatal kinetics in either species, while revealing modest early stage C_4_-specific adjustments in stomatal dynamics without affecting photosynthetic capacity or biochemical parameters.
- We demonstrate that cereal grafting does not alter core physiological traits in rice or pearl millet and can be deployed without long-term impact on photosynthesis. These findings establish cereal grafting as a tractable platform for mechanistic dissection of root-shoot signaling and trait combination across different C_3_ and C_4_ cereals.

## Introduction

Grafting is an ancient horticultural practice in which a shoot (scion) and a root system (rootstock) from two different plants are joined, establishing vascular continuity and enabling growth as a single composite plant (Melnyk & Meyerowitz, 2015; Gautier *et al*., 2019; Kragler & Bock, 2025). The scion and rootstocks may be derived from either the same or different species, depending on the intended application and compatibility between the donor tissues (Xiao *et al*., 2023; Kragler & Bock, 2025). For centuries, grafting has been used primarily in dicotyledonous crops (Mudge *et al*., 2009; Kragler & Bock, 2025). In contrast, grafting in cereal crops was long considered impossible because monocotyledonous species lack a vascular cambium and possess scattered vascular bundles, anatomical features thought to preclude successful graft union formation (Muzik & La Rue, 1952; Gautier *et al*., 2019; Rasool *et al*., 2020). This long-standing consensus was recently overturned by the successful grafting of cereals at the embryonic root-shoot interface, establishing functional vascular continuity between root and shoot systems (Reeves *et al*., 2022). The developmental plasticity of embryonic tissues, including the meristematic activity of the mesocotyl, enables the formation of *de novo* vascular connections integrating root and shoot systems despite the absence of a vascular cambium in cereals (Gaillochet & Lohmann, 2015; Reeves *et al*., 2022). Although the cellular and molecular mechanisms underlying graft union formation in cereals remain poorly understood, the capacity for vascular neogenesis distinguishes embryo-dependent graft-union formation in cereals from the cambium-dependent process that underpins grafting in dicotyledonous species (Mukundan *et al*., 2024). This establishes the use of embryonic tissues for cereal grafting as both a functional tool in cereal biology and a promising experimental platform for trait combination across species.

Previous studies on grafting in dicotyledonous crops demonstrate influence in broad range of photosynthesis-related processes. For example, grafting cultivated tomato (*Solanum lycopersicum*) scions onto wild relative rootstocks (*Solanum pennellii*) influences stomatal development and density, changes that led to high photosynthetic assimilation in grafted tomato compared to non-grafted (Ganie *et al*., 2025). In another study, grafting muskmelon on two different inter-specific rootstocks (*Cucurbita maxima* x *C. moschata*) increased net photosynthesis rate, stomatal conductance, concentration of intercellular CO_2_ and transpiration rate (Liu *et al*., 2011). In the perennial woody species grapevine (*Vitis vinifera*), rootstocks modulate scion stomatal conductance and transpiration by regulating the water transport capacity of the root system (Alsina *et al*., 2011; Marguerit *et al*., 2012). Further examples and information on grafting-mediated effects on plant growth and physiology in dicots have been reviewed extensively (Fullana-Pericàs *et al*., 2020; Kragler & Bock, 2025). The physiological shifts could arise from the grafting process itself, which involves callus formation, vascular reconnection and the re-establishment of symplastic and apoplastic continuity, potentially inducing transient or persistent changes in carbon allocation and stomatal behavior (Melnyk, 2017; Wang *et al*., 2024; Kragler & Bock, 2025). Whereas, grafting in dicotyledonous species is typically performed on established plants with differentiated tissues, cereal grafting is performed at the seed stage, capitalizing on the meristematic activity of embryonic mesocotyl tissue to enable graft union formation (Reeves *et al*., 2022). Nevertheless, the physiological consequences of grafting in cereals remain largely unexplored, and whether the grafting process itself alters photosynthetic performance or stomatal dynamics has not been determined.

Rice (*Oryza sativa*) and pearl millet (*Cenchrus americanus*, syn. *Pennisetum glaucum*) are phylogenetically divergent cereals belonging to the BEP and PACMAD clades of the Poaceae, respectively, and differ fundamentally in photosynthetic biochemistry (Pardo & VanBuren, 2021; Wilson & VanBuren, 2022). Rice utilizes C_3_ photosynthesis, in which the dual carboxylase/oxygenase activity of Rubisco renders carbon assimilation sensitive to photorespiration, whereas pearl millet employs a C_4_ CO_2_-concentrating mechanism that suppresses photorespiration and confers greater nitrogen- and water-use efficiency (Ghannoum *et al*., 2010; Pardo & VanBuren, 2021; Wilson & VanBuren, 2022). This phylogenetic and physiological breadth makes them valuable comparative models for evaluating whether cereal grafting affects physiological traits across divergent grass lineages. Moreover, under natural field conditions, plants experience frequent and rapid fluctuations in light that require coordinated stomatal and biochemical adjustments (Acevedo-Siaca & McAusland, 2025). Water availability similarly shapes photosynthetic performance, and rice and pearl millet occupy contrasting ecological water niches (Bouman, 2009; Serba *et al*., 2020). Whether cereal grafting alters steady-state photosynthesis or dynamic responses to light under contrasting water regimes and across developmental timepoints has not been examined. By comparing grafted and non-grafted rice and pearl millet under contrasting water regimes and at different plant developmental stages, our study tests whether cereal grafting imposes physiological consequences on carbon gain, water-use and stomatal-photosynthetic coordination, and whether any such effects are conserved across divergent photosynthetic strategies or are species-specific.

Our study provides the first comprehensive physiological assessment of cereal grafting across photosynthetically distinct C_3_ and C_4_ species. Despite fundamental differences in photosynthetic metabolism, stomatal regulation and water-use strategies between rice and pearl millet, cereal grafting imposed no substantial physiological constraints on plant performance. In both species, plant growth, photosynthetic capacity, stomatal kinetics and stomatal-photosynthetic coordination were largely maintained across all graft combinations and water regimes. This demonstrates that cereal grafting preserves the core functional characteristics of non-grafted plants. Although pearl millet exhibited modest early-stage stomatal adjustments, these responses were transient and were not associated with long-term changes in photosynthetic biochemistry, carbon assimilation or water-use efficiency.

## Materials and Methods

### Plant material and cereal grafting

Mature seeds of rice (*Oryza sativa* japonica var. Kitaake) and pearl millet (*Cenchrus americanus*, syn. *Pennisetum glaucum*) were surface sterilized in 35 % (v/v) sodium hypochlorite for five minutes at room temperature, and then rinsed thoroughly with sterile Milli-Q water. The seeds were transferred to 50 ml Falcon tubes containing few drops of water and imbibed in darkness at 28 °C for 16 hours in an incubator (Binder, BINDER GmbH, Tuttlingen, Germany). We grafted the imbibed seeds following procedures previously described by (Reeves *et al*., 2022) and protocols optimized in our laboratory. We generated three graft combinations, (i) non-grafted controls (NG), (ii) self-grafts (SG) in which embryonic scion was excised and reattached to the same seed, and (iii) intra-species grafts (IG) in which embryonic scion was excised and exchanged with the embryonic scion of a different seed of the same species. After grafting, the grafted-seeds were transferred onto sterile petri plates lined with two moist Whatman filter papers (Grade 1, Whatman plc, Maidstone, United Kingdom) and incubated in a Conviron GEN1000-ECO plant growth chamber (Conviron, North Dakota, USA) under controlled conditions: 27 / 23 °C (day / night), 65 % relative humidity and a 12-hour photoperiod. We provided photosynthetically active radiation (PAR) of 400 μmol m^−2^ s^−1^ using neutral-white T5 LED lamps (model T5-900EC-12W, 12W, Shenzhen, China). Grafted seeds were monitored daily, with moisture maintained as required until both root and shoot emerged. After seven days, successful grafts were selected following previously described procedures (Reeves *et al*., 2022) and established laboratory protocols, transplanted and used for subsequent experiments.

### Plants growth conditions and experimental design

We transplanted seven days old rice and pearl millet seedlings into 1L pots filled with non-sterilized potting substrate (Levington’s Advance F2+S compost, medium nutrient). We maintained the seedlings under controlled water regime by irrigating pots to 160 % of the substrate’s field water holding capacity (FC). To determine the FC level, we followed the gravimetric procedure established by Cassel & Nielsen (1986). Briefly, we saturated pots containing substrate with water, allowed them to drain freely, and recorded the saturated weight. We then oven-dried the substrate at 80 °C for 72 hours, and recorded the dry weight. We calculated amount of moisture equal to 100 % FC as the difference between saturated and dry weights. We grew plants in a controlled environment chamber (Aralab, Portugal) which is part of the Smart Technology Experimental Plant Suite (STEPS) at the School of Life Sciences, University of Essex, United Kingdom. We maintained the non-grafted controls at the same growth conditions as those used for self- and intra-species grafts. Aralab plant growth chambers operated with broad-spectrum white LED lamps for radiation (model VR-3p-BW4, Austin, USA). We manually weighed each pot and subsequently calculated the volume of water required to maintain 160 % FC until the plants were 35 days old, counting from the day of seed grafting. At 35 days after grafting (DAG), we established a fully factorial experiment with two water regimes; 160 % FC (hereafter saturated) and 100 % FC (hereafter optimal), and two sampling timepoints; timepoint-1 at 38 DAG (three days after withholding irrigation, and onset of optimal water regime), and timepoint-2 at 43 DAG (five days of sustained irrigation under option water regime) (Fig. **1a**). At each experimental timepoint, we determined steady-state photosynthetic performance by measuring gas-exchange and dynamic stomatal responses (see gas exchange measurement section below) (Fig. **1a**). We included a minimum of three biological replicates per graft type within each water regime and timepoint. Throughout the experiment, each plant was supplemented weekly with 100 mL of nutrient solution containing 0.34 g L^−1^ of Solufeed Fe 13.2 % EDTA (Solufeed Ltd., United Kingdom) and 0.065 g L^−1^ of Peters Excel CalMag Grower (ICL Growing Solutions, ICL Group, United Kingdom).

**Fig 1.**
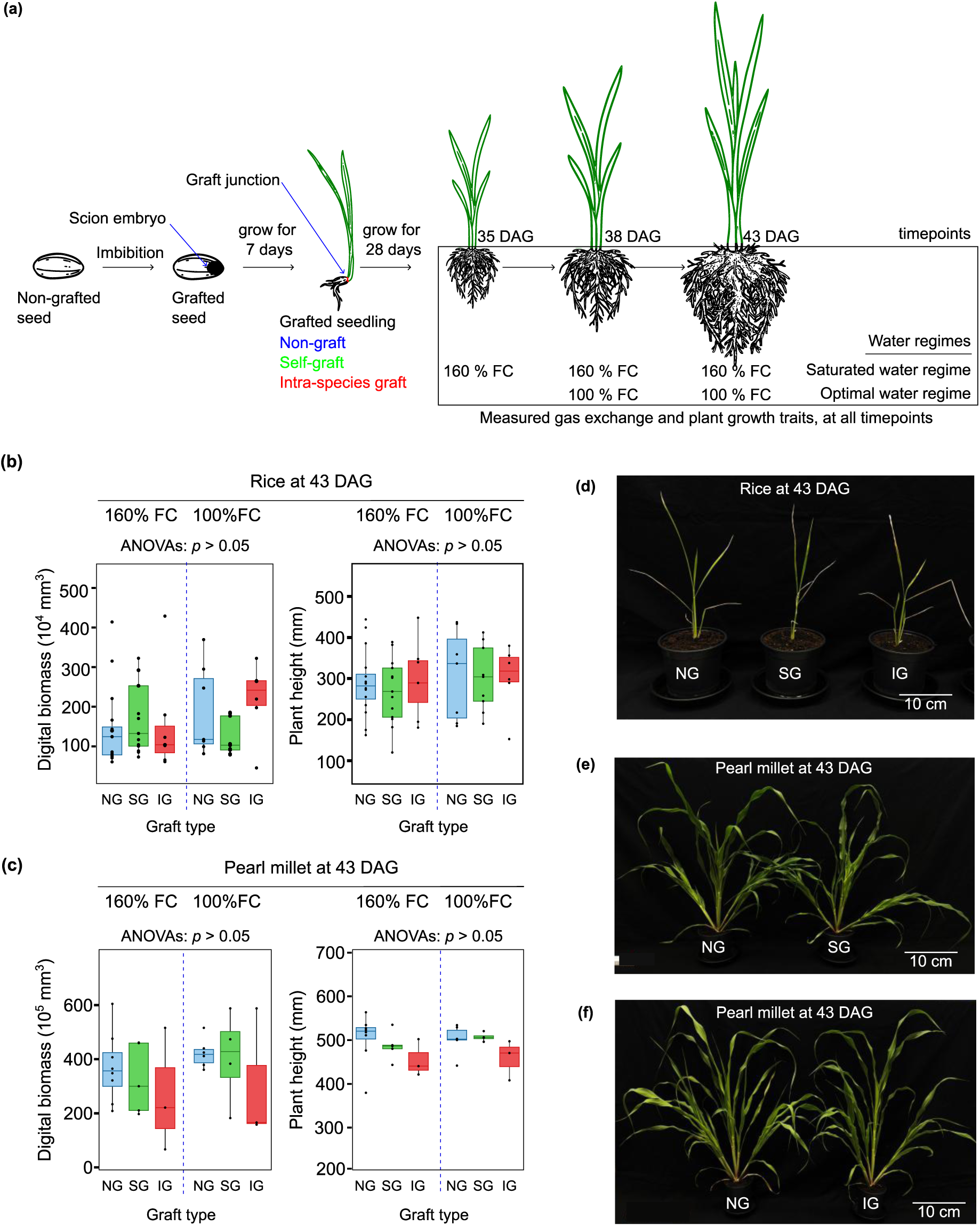
Experimental design and phenotypic responses of rice and pearl millet plants under contrasting water regimes. **(a) Schematic representation of the experimental design:** Mature rice and pearl millet seeds were used for cereal grafting to generate three combinations: non-grafts (NG), self-grafts (SG) and intra-species grafts (IG). The grafted plants were grown under saturated water regime of 160 % of soil water field capacity (FC) until 35 days after grafting (DAG). An additional optimal water regime (100 % FC) was imposed thereafter. Plants required three days to reach optimal water regime (timepoint 1, 38 DAG) and were maintained at this soil-moisture level for a further five days (timepoint 2, 43 DAG). Control plants under the saturated water regime were maintained throughout. At each timepoint, physiological performance was assessed using both steady-state and dynamic gas-exchange measurements and plant growth traits were recorded. Panels **(b)** and **(c)** are **growth measurements of rice and pearl millet plants at 43 DAG**: Plant digital biomass and height were measured across different graft types of rice and pearl millet plants. Panels **d-f** are representative images of rice and pearl millet graft types. Blue, green and red colors represent NG, SG and IG, respectively. Data of time point 3 (43 DAG) is presented as mean ± SEM (*n* = 3 - 16). Two-way ANOVA was used to evaluate the main effect of graft type and water regime, and their interaction.

### Phenotypic measurements

We measured plant growth traits, using the PlantEye F600 multispectral 3D scanner (Phenospex, The Netherlands). The PlantEye is ideal for plant phenotyping because it is non-destructive and can provide high-resolution and real-time data on plant growth and health by combining 3D and multispectral imaging. We measured the plant growth traits at each experimental timepoint. For each trait and timepoint, we captured three replicate multispectral images per plant and averaged the values to obtain a single representative measurement value.

### Gas exchange measurements

We conducted gas exchange measurements, on the youngest fully expanded leaf of each plant. These measurements included (i) photosynthetic responses to sequential step changes in light intensity and quality, and (ii) net assimilation in response to intercellular CO_2_ (*A/Ci* curves). We performed the measurement using the LI-COR 6800 (LI-COR Biosciences Inc., Nebraska, USA) equipped with a 6800-01A Multiphase Flash Fluorometer chamber (MPF, LI-COR). We set the environmental cuvette conditions to 27 °C, 400 µmol mol^−1^ CO_2_ and 65 % relative humidity. The MPF system supplied actinic light using a variable red:blue light ratio depending on the measurement type. At each timepoint, we first performed the light step-responses, prior to *A/Ci* curves, to optimize experimental time. All gas exchange measurements were completed before 14:00 to minimize potential diurnal or circadian effects on photosynthetic measurements.

#### (i) Photosynthetic *A/Ci* response curves

We assessed the net assimilation rate in response to intercellular CO_2_ (*A/Ci*) curves. We measured and recorded several parameters, including stomatal conductance (*g_sw_*), net photosynthetic CO_2_ assimilation rate (*A*) and intercellular CO_2_ (*Ci*) according to von Caemmerer & Farquhar (1981). We calculated the intrinsic water-use efficiency (*iWUE*) as *A* divided by *g_sw_* (*A*/*g_sw_*). To characterize photosynthetic performance, we generated *A/Ci* curves for rice and pearl millet under the same environmental conditions, with actinic light set to 1500 µmol m^−2^ s^−1^ at a 90:10 red:blue light ratio. Once each leaf reached steady-state at 400 µmol mol^−1^ CO_2_, we measured gas exchange in rice using the following reference CO_2_ concentrations: 400, 300, 200, 100, 50, 400, 550, 700, 900, 1200, 1500, 1800 and 2000 µmol mol^−1^. In the case of pearl millet, the reference CO_2_ concentrations were 1250 (twice to ensure stabilization),1000, 800, 600, 400, 350, 300, 250, 200, 150, 100, 65, 40, 20, 10 and 0 µmol mol^−1^. We logged gas-exchange values every 150 seconds at each CO_2_ concentration level. We used the resulting CO_2_ curves to derive the core photosynthetic parameters at 400 µmol mol^−1^ point in the *A/Ci* curves. From the gas exchange data we estimated core photosynthetic parameters by fitting mechanistic models to the *A/Ci* curves using the PhotoGEA R package (Lochocki *et al*., 2025). For rice which is a C_3_ crop, we fitted the Farquhar-von Caemmerer-Berry (FvCB) model (Farquhar *et al*., 1980) and applied the temperature functions described by Sharkey and Bernacchi to normalize *V_c.max_* (maximum carboxylation rate) and *J_max_* (maximum electron transport rate) at 25 °C (Bernacchi *et al*., 2001; Sharkey *et al*., 2007). For pearl millet which is a C_4_ crop, we fitted the C_4_-specific mechanistic model described by von Caemmerer (2000). To assess the model stability across the pearl millet measurements, we estimated *V_c.max_* and *V_p.max_* (maximum PEP carboxylation rate) while fixing the *Vs_pr_* (PEP-regeneration rate) and *J* (electron-transport rate) at 1000 µmol m^−2^ s^−1^. We also determined *A_max_* (maximum rate of net CO_2_ assimilation rate) for both plant species as the light-saturated assimilation rate reached the highest CO_2_ concentrations.

#### (ii) Stomatal induction kinetics

We measured the photosynthetic induction in response to step changes in light intensity and quality, using dark-acclimated leaves. We first applied 100 µmol m^−2^ s^−1^ actinic light (100 % red light) and allowed *A* and *g_sw_* to stabilize until they reached steady-state values. Stabilization required between 15 - 20 minutes for rice and 30 - 35 minutes for pearl millet. We maintained the 100 µmol m^−2^ s^−1^ red light for 15 minutes, before increasing the light intensity in a single step to 1500 µmol m^−2^ s^−1^ of 100 % red light for 30 minutes. Finally, we adjusted the spectrum to percentage ratio of 90:10 red:blue light at 1500 µmol m^−2^ s^−1^ for an additional 15 minutes. We logged gas exchange data every 5 seconds throughout the induction sequence. To quantify the induction speed for both photosynthesis and stomatal responses, we fitted the two-parameter Cumulative Distribution Weibull (CDWeibull) model to the gas exchange data *A* and *g_sw_* (Weibull, 1939; Woning *et al*., 2026). This model allowed us to capture exponential and sigmoidal responses without imposing a pre-defined curve assumptions (Woning *et al*., 2026). We modelled the *A* and *g_sw_* response in our plants over experimental time using the following equation:

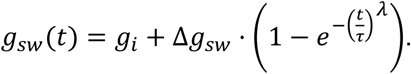

In this model, *g_i_* represents the initial steady-state value of *g_sw_* value before light step, and *Δg_sw_* represents the total variation between the initial and final steady-state values. The parameter τ (tau) represents the time constant required to achieve approximately 63 % of this total variation, while λ (lambda) is a unitless constant that determines the curvature or initial lag of the response. In this study, we extracted *τ_A_* and *τ_gsw_* as the primary estimators of induction speed. A smaller *τ* value indicates fast response to the changes in light, whereas a larger value suggests delayed response.

### Statistical analyses

Data were analyzed using open source software RStudio in R environment (Posit team, 2025; R Core Team, 2025). Both plant growth and core photosynthetic parameters derived at 400 µmol m^−1^ in the *A/Ci* curves processed using suite of libraries in the R package tidyverse (Wickham *et al*., 2019). We performed one-way ANOVA for all plant growth and photosynthetic parameters derived at 400 µmol m^−1^ in the *A/Ci* curves data sets recorded at 35 DAG, and two-way ANOVA for all similar data sets recorded at the timepoint-1 (38 DAG) and timepoint-2, (43 DAG), to determine the main effects emerging from cereal grafting alone, water regimes alone, or their interactions, across the experimental timepoints. In cases where the ANOVA results were significant, the differences between the experimental groups per timepoint were detected using Tukey’s Honest Significant Difference (HSD) for multiple comparisons at *p* ≤ 0.05. In all cases, the data were visualized using the R package ggplot2 (Wickham, 2016).

## Results

### Cereal grafting does not alter growth traits in rice and pearl millet under contrasting water regimes

To determine the physiological consequences of cereal grafting, we evaluated plant growth traits in the two photosynthetically distinct species: rice and pearl millet. We focused on plant biomass and height across different graft combinations, under saturated and optimal water regimes (Fig. **1**; Table **S1**, **S2**). At 38 DAG, corresponding to onset of optimal water regime, we observed marginal temporary shift in biomass (Table. **S1**; see main effect of interaction of graft x regime, *p* = 0.062), but this effect was not statistically significant and did not persist upon sustained optimal water regime corresponding to 43 DAG (Fig. **1b**, Table. **S1**; see 43 DAG). At the end of the experiment, rice plants exhibited similar growth across all graft combinations irrespective of the water regime (Fig. **1d**). Our data indicate that in rice neither graft type nor water regime significantly influenced biomass or height across the experimental timepoints (Table. **S1**). In pearl millet, grafting had stronger influence on the growth traits. Biomass showed marginal but statistically non-significant response to graft type at saturated water regime (Table. **S2**; see 35 DAG; main effect of grafting *p* = 0.065) and onset of optimal water regime (Table. **S2**; see 38 DAG; main effect of grafting *p* = 0.074). In contrast, height was significantly affected by grafting at the same timepoints (Table. **S2**; see 35 DAG, main effect of grafting *p* = 0.018; and at 38 DAG, main effect of grafting *p* = 0.038). We further observed that these grafting effects diminished upon sustained optimal water regime, suggesting the plants fully stabilized (Fig. **1c**, **e, f**; Table. **S2**; see 43 DAG). Across both species, water regime alone or the interaction between graft type and water regime did not affect the growth traits (Table. **S1**, **S2**). These results indicate that while cereal grafting and the transition to an optimal water regime weakly modulates early growth trajectories, particularly in pearl millet, grafted plants ultimately achieved growth trends comparable to non-grafts across both watering regimes.

To better understand the effects of cereal grafting on core plant physiological traits and to interpret the observed growth responses, we first assessed the impact of cereal grafting on steady-state and dynamic photosynthetic performance in rice and pearl millet under saturated conditions at 35 DAG. We then evaluated how these responses changed over time, including the transition to an optimal water regime at 38 DAG and following sustained optimal watering at 43 DAG.

### Cereal grafting induces early-stage, C_4_ species-specific stomatal adjustments without affecting photosynthetic biochemistry

We evaluated net photosynthetic CO_2_ assimilation rate (*A*) in response to intercellular CO_2_ concentration (*Ci*) in grafted rice and pearl millet under a saturated water regime (Fig. **2**). In rice, we observed similar photosynthetic performance across all graft combinations (Fig. **2a**).

**Fig. 2.**
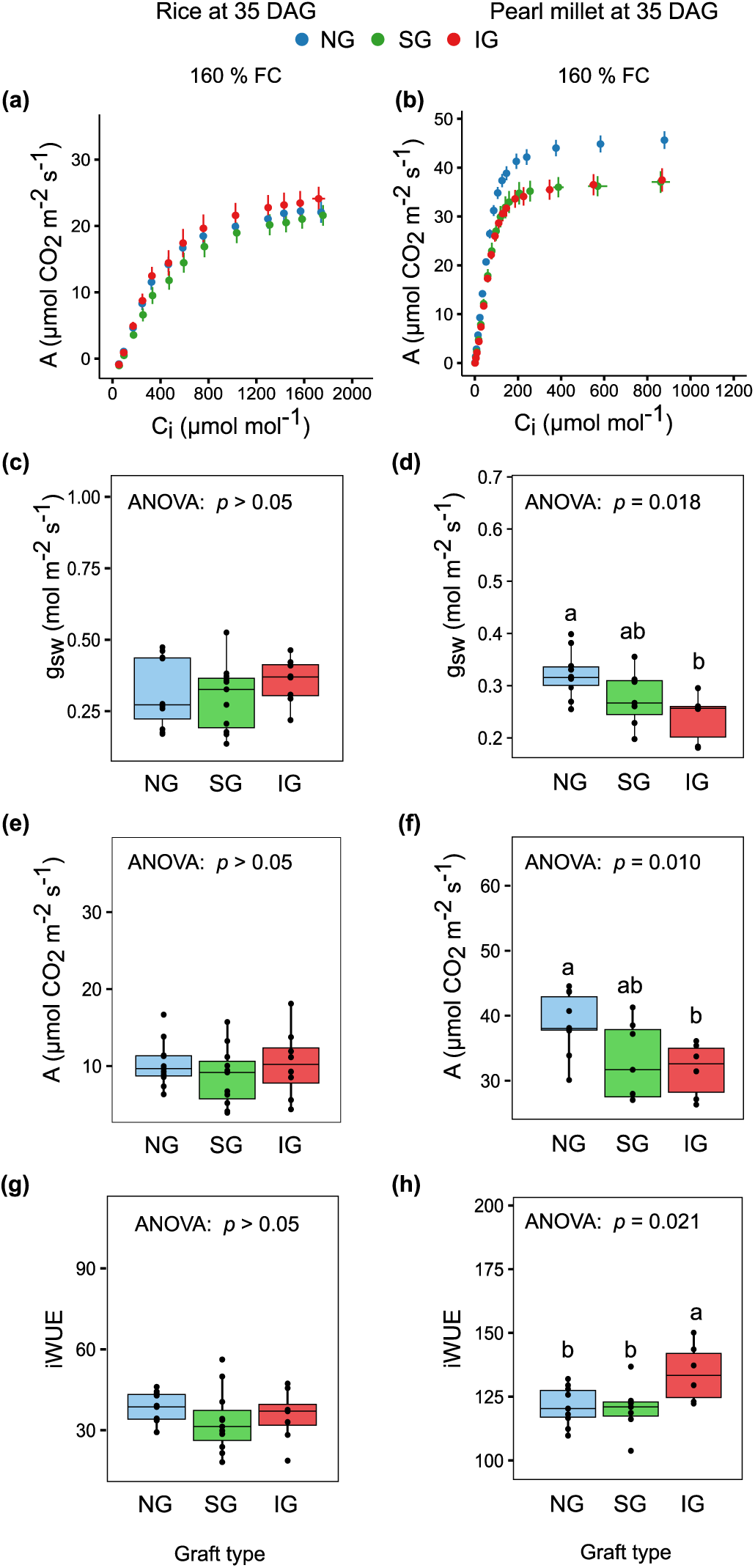
Gas exchange measurements and derived photosynthetic parameters of in rice and pearl millet. CO_2_ response curves were measured in rice and pearl millet non-grafts (NG), self-grafts (SG) and intra-species grafts (IG) at saturated water regime of 160 % of soil water field capacity at 35 days after grafting. Panels **(a, b)** shows Net photosynthetic assimilation (*A*) as a function of intercellular CO_2_ (*Ci*) *A/Ci* curve in rice (50 - 2000 µmol mol^−1^) and pearl millet (0- 1250 µmol mol^−1^), respectively. Panels **(c, d)** shows stomatal conductance (*g_sw_*) at 400 µmol mol^−1^ in rice and pearl millet, respectively. Panels **(e, f)** shows *A* at 400 µmol mol^−1^ in rice and pearl millet, respectively; and panels **(g, h)** shows intrinsic water use efficiency (*iWUE*) at 400 µmol mol^−1^ in rice and pearl millet, respectively. Blue, green and red colors represent NG, SG and IG, respectively. Data in panels **c** - **h** is presented as mean ± SEM (*n* = 6 - 11). One-way ANOVA was used to evaluate the effect of graft type, and where applicable statistical differences were assessed using Tukey’s HSD post hoc test.

Likewise, there were no statistically significant differences in key photosynthesis parameters, including *V_c,max_*, *J_max_* and *A_max_* (Table. **1** and Table **S3;** see rice). Moreover, *g_sw_*, *A* and *iWUE* derived at 400 µmol mol^−1^ CO_2_ showed no significant differences across the graft combinations (Fig. **2c, e, g**, Table. **S4**; see rice). In pearl millet, the non-grafts exhibited higher photosynthetic rates compared to self- and intra-species grafts (Fig. **2b**). Concomitantly, non-grafts showed significantly higher *A_max_* values (∼ 46 µmol m^−2^ s^−1^) compared to values in both self and intra-species grafts (∼ 38 µmol m^−2^ s^−1^) (Table. **1**; Table. **S3;** see pearl millet). There were no differences in *V_c,max_* and *V_p,max_* between grafted and non-grafted pearl millet (Table. **1** and Table. **S3**; see pearl millet). Moreover, we observed reduced *g_sw_* and *A* in self- and intra-species grafts compared to non-grafts, although this effect was only statistically significant in intra-species grafts (Fig. **2d, f**; Table. **S4**; see pearl millet). Consequently, the leaf-level water-use efficiency was higher in self- and intra-species pearl millet grafts, although the increase was only statistically significant in intra-species grafts (Fig. **2h**; Table. **S4**; see pearl millet).

**Table 1.**
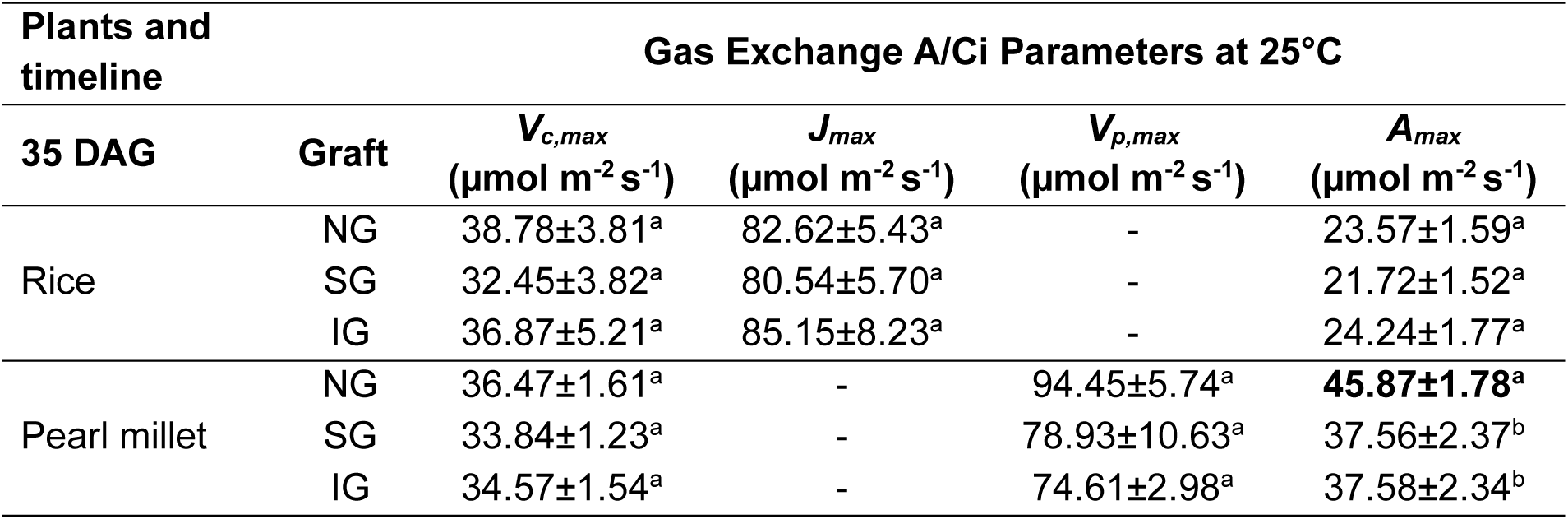
Key photosynthetic parameters of rice and pearl millet under saturated water regime. Key photosynthesis parameters, including maximum carboxylation rate (*V_c,max_*), maximum assimilation rate (*A_max_*), maximum rate of electron transport (*J_max_*) and maximum PEP Carboxylase carboxylation rate (*V_p,max_*) were derived from CO_2_ response curves generated at 25 °C in rice and pearl millet, non-grafts (NG), self-grafts (SG) and intra-species grafts (IG), under saturated water regime of 160 % of soil water field capacity at 35 days after grafting. We derive estimates of the maximum Rubisco carboxylation rate (*V_c,max_*) and maximum assimilation rate (*A_max_*) in both plants. While maximum rate of electron transport (*J_max_*) was estimated in rice only, and the maximum PEP Carboxylase carboxylation rate (*V_p,max_*) in pearl millet. Data are means ± SEM (*n* = 6 - 11). One-way ANOVA was used to evaluate the effect of graft type on the photosynthesis parameter. Different lowercase superscript letters within columns indicate significant differences (*p* < 0.05) between the graft types as determined by Tukey’s HSD post-hoc test.

Together, these early-stage responses in pearl millet suggest a modest stomatal adjustment following grafting. However, the stability of core carboxylation parameters (*V_c,max_* and *V_p,max_*) in both species indicates that cereal grafting does not impair the biochemical capacity for photosynthesis. These findings suggest a transient physiological adjustment rather than a sustained disruption of photosynthetic function, and further indicate that the observed responses may be specific to the photosynthetic type rather than the grafting process per se.

### Cereal grafting does not alter dynamic stomatal kinetics but reveals C_3_-C_4_ specific differences in stomatal-photosynthetic coordination

To understand the impact of cereal grafting on rice and pearl millet stomatal traits, we first evaluated *g_sw_* responses of grafted plants to step changes in light intensity and quality (Fig. **3a, b**), as photosynthetic performance and gas exchange are strongly influenced by the dynamic response of stomata to light. This step change approach allowed us to distinguish stomatal aperture responses driven by photosynthetic activity (red light) from those regulated by phototropin-mediated signaling pathways (blue light). The *g_sw_* in rice self- and intra-species grafts showed no substantial differences compared to non-grafts (Fig. **3a**). In contrast, non-grafted pearl millet exhibited higher *g_sw_* under step changes in light intensity and quality (Fig. **3b**). This increased stomatal conductance may account for the higher *A* and *A_max_* observed previously (see Fig. **2b**; Table. **1**; see pearl millet), because enhanced CO_2_ diffusion supports higher carboxylation rates under increasing intercellular CO_2_ availability.

**Fig 3.**
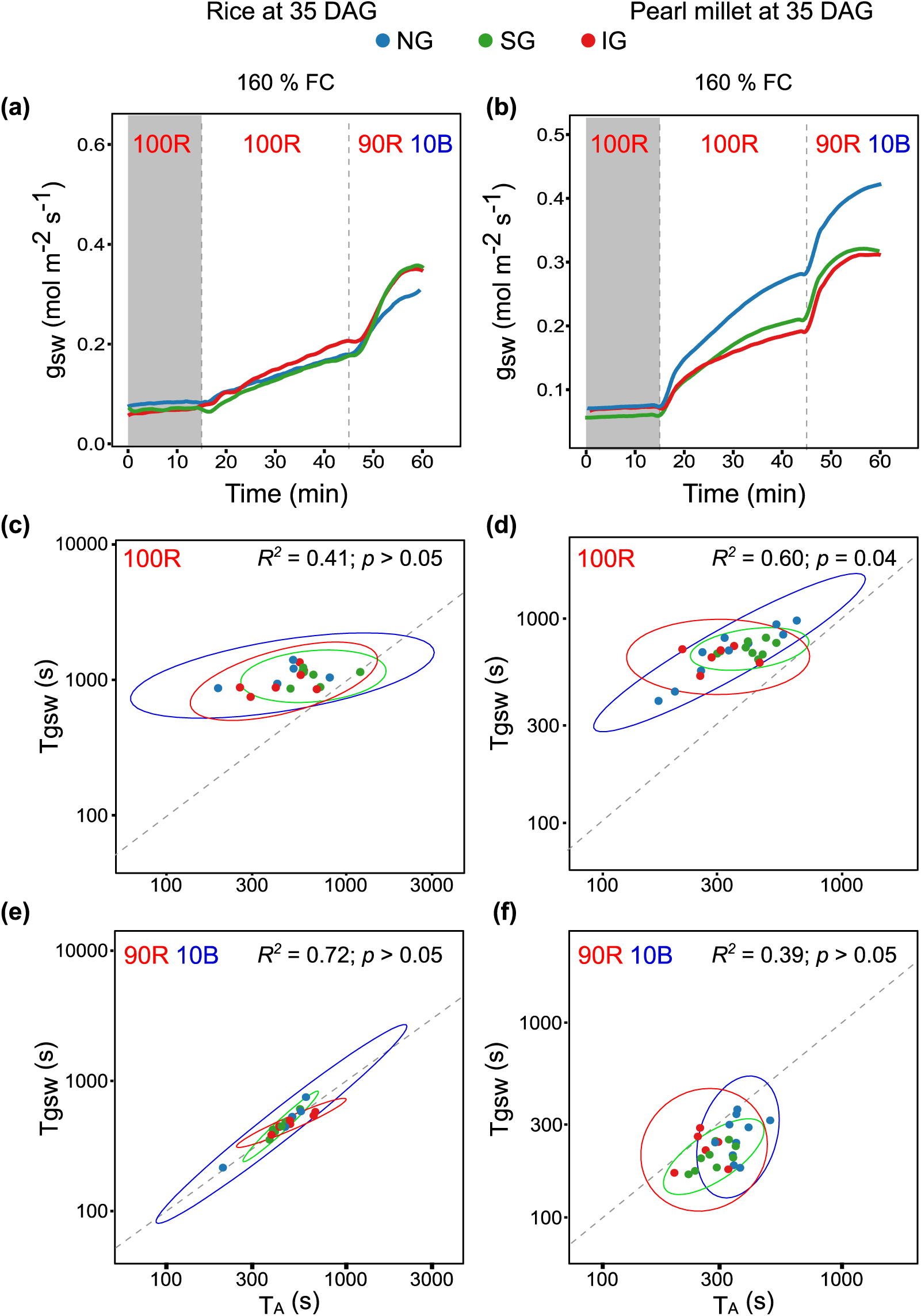
Dynamic photosynthetic and stomatal responses to light induction in rice and pearl millet. The effects of light intensity and quality on stomatal kinetics were assessed in both rice and pearl millet, non-grafts (NG), self-grafts (SG) and intra-species grafts (IG) grown at saturated water regime of 160 % of soil water field capacity at 35 days after grafting (DAG). Panels **(a**, **b)** show stomatal conductance (*g_sw_*) response under sequential step-changes in light intensity and quality in rice and pearl millet, respectively. The regions in grey color represent periods of light transition from low light (100 µmol m^−2^ s^−1^) to high light (1500 µmol m^−2^ s^−1^). Variation of light quality was as follows: 100 µmol m^−2^ s^−1^ red light for 15 min, to 1500 µmol m^−2^ s^−1^ red light for 30 min and 1500 µmol m^−2^ s^−1^ and 90:10 red:blue light ratio for 15 min. Panels **(c - f)** are modelled correlations between the time constants of stomatal conductance (τ_gsw_) and net photosynthetic CO_2_ assimilation (τA), under saturated water regime, 35 DAG. Panels **(c, d)** show the responses during the transition to high red light (1500 µmol m^−2^ s^−1^), while panels **(e, f)** show the responses during subsequent transitions to 1500 µmol m^−2^ s^−1^ at a 90:10 red:blue light ratio. Time constants represent the time required to reach 63 % of the steady-state maximum for each parameter following the light transitions. The dashed line represents the 1:1 coordination line between τ_gsw_ and τA. In panel **c-f**, the blue, green and red lines represent NG, SG and IG, respectively. Data in panel **a** and **b** are presented as mean ± SEM (*n* = 5 - 10).

Next, we quantified the dynamic coordination between *g_sw_* and *A*, by calculating the time constants (τ) required to reach 63 % of the steady-state maximum for both parameters following step changes in light intensity and quality (Fig. **3c-f**; Tables. **S5**). Under red light, average *τ_gsw_* and *τ_A_* in rice were on average about 17 min and 9 min respectively, and both decreasing to, on average about 8 min under blue light (Table. **S5**; see rice). Across all transitions, no significant differences in *τ_gsw_* or *τ_A_* were detected among grafted rice, indicating that grafting did not alter stomatal or photosynthetic induction kinetics (Table. **S6**; see rice). In pearl millet, the average *τ_gsw_* and *τ_A_* under red light were on average 11 min and 6 min (Table. **S5**; see pearl millet red light). Under blue light, self- and intra-species pearl millet grafts exhibited significantly faster *τ_gsw_* and *τA*, on average about 4 min and 3 min, respectively, compared to non-grafted controls, which had 6 min and 4 min, *τ_gsw_* and *τA*, respectively (Table. **S5, S6**; see pearl millet, blue light). These results suggest that the lower *g_sw_* of grafted plants may enable them to reach steady-state values more rapidly during light transitions, likely due to a smaller absolute change required to adapt compared to non-grafts. In both species and across all graft types, the initial transition to high red light irradiance suggested that photosynthetic induction may be partly constrained by slower stomatal opening (Fig. **3c, d**). In contrast, the addition of 10 % blue light reduced the lag between *τ_gsw_* and *τ_A_* (Fig. **3e, f**). In rice, *τ_gsw_* and *τ_A_* converged towards a 1:1 relationship (Fig. **3e**; Table. **S7**; see rice, blue light), while pearl millet displayed a distinct pattern (Fig. **3f**; Table. **S7**; see pearl millet, blue light), where *τ_gsw_* was consistently lower than *τA*, with data points falling below the 1:1 coordination line. Importantly, these species-specific coordination patterns were conserved across the graft combination, indicating that cereal grafting does not alter the underlying signaling that coordinates stomatal dynamics with photosynthetic induction. Instead, the observed differences reflect inherent C_3_ and C_4_ photosynthetic type dependent differences in the regulation of stomatal-photosynthetic coupling during dynamic light responses.

### Steady-state photosynthetic performance remains stable under sustained optimal water regime following graft establishment

After establishing that cereal grafting had minimal to no effects on both steady-state and dynamic physiological performance under saturated water regime, we investigated whether these responses were maintained under optimal water regime over a sustained 5-day period and across a longer developmental timeframe (Fig. **1a**). This enabled us to determine whether the reduced *g_sw_* and *A* previously observed in pearl millet at 35 DAG persisted during later stages of growth and varying water regimes. Consistent with observations at 35 DAG, photosynthetic performance remained similar across all rice graft combinations under saturated water regime, onset of optimal water regime and when sustained for 5-day period (Fig. **4a, b**). Likewise, we found no differences in photosynthesis parameters, including *V_c,max_, J_max_* and *A_max_* across all respective rice graft combinations and water regime (Table. **S8**; see rice). Although changes in photosynthetic parameters *V_c,max_, J_max_* and *A_max_* were not statistically significant, values tended to increase with plant age. Between 38 DAG and 43 DAG *V_c,max_* increased on average by 14 µmol m^−2^ s^−1^, *J_max_* by 20 µmol m^−2^ s^−1^ and *A_max_* by 3 µmol m^−2^ s^−1^ (Table. **2**; Table. **S8**; see rice at 38 and 43 DAG). This stability in photosynthesis was reflected in *g_sw_*, *A* and *iWUE* (Fig. **4c-h**; Table. **S9**; see rice). In pearl millet, at both 38 and 43 DAG, all graft combinations displayed comparable *A/Ci* responses (Fig. **5a, b**). In contrast, derived photosynthetic parameters showed variation driven by significant main effects of grafting and water regimes (Table. **S8**; see pearl millet). At 38DAG, we observed graft mediated increase in *V_c,max_, V_pmax_* alongside a water regime mediated increase in *A_max_* by on average 5 µmol m^−2^ s^−1^ under optimal compared with saturated water regime (Table. **2**; Table. **S8**; see pearl millet 38 DAG). However, at 43 DAG, corresponding to 5-day sustained optimal water regime, these differences were no longer evident. Mean *V_c,max_* was 28 µmol m^−2^ s^−1^ under optimal water condition compared to 29 µmol m^−2^ s^−1^ under saturated water regime, while *A_max_* was 37 µmol m^−2^s^−1^ under optimal water regime compared to 36 µmol m^−2^ s^−1^ under the saturated water regime (Table. **2**; see pearl millet 43 DAG; Table. **S8**; see pearl millet). Only *V_pmax_* remained elevated under optimal water regime (108 µmol m^−2^ s^−1^) compared to under saturated water regime (92 µmol m^−2^ s^−1^) consistent with a marginal non-significant effect of water regime (Table. **S8**; see pearl millet *V_pmax_* regime *p* = 0.059). Likewise, shifts in *g_sw_*, *A* and *iWUE* were evident at 38 DAG (Fig. **5c, e, g**; Table. **S9**; see pearl millet), but were no longer detectable at 43 DAG (Fig. **5d, f, h**; Table. **S9**; see pearl millet). At 38 DAG, changes in *A* and *g_sw_* were primarily driven by water regime (Table. **S9**; see pearl millet 38 DAG, regime (*A, gsw*): *p* = 0.035, *p* = 0.036, respectively), whereas *iWUE* was influenced by interaction between graft type and water regime (Table. **S9**; see pearl millet 38 DAG, graft x regime (*iWUE*): *p* = 0.032). Collectively, these results indicate that the differences in gas exchange and photosynthetic capacity observed in pearl millet at 35 DAG weakly persist at 38 DAG but dissipate during later developmental stages.

**Fig. 4.**
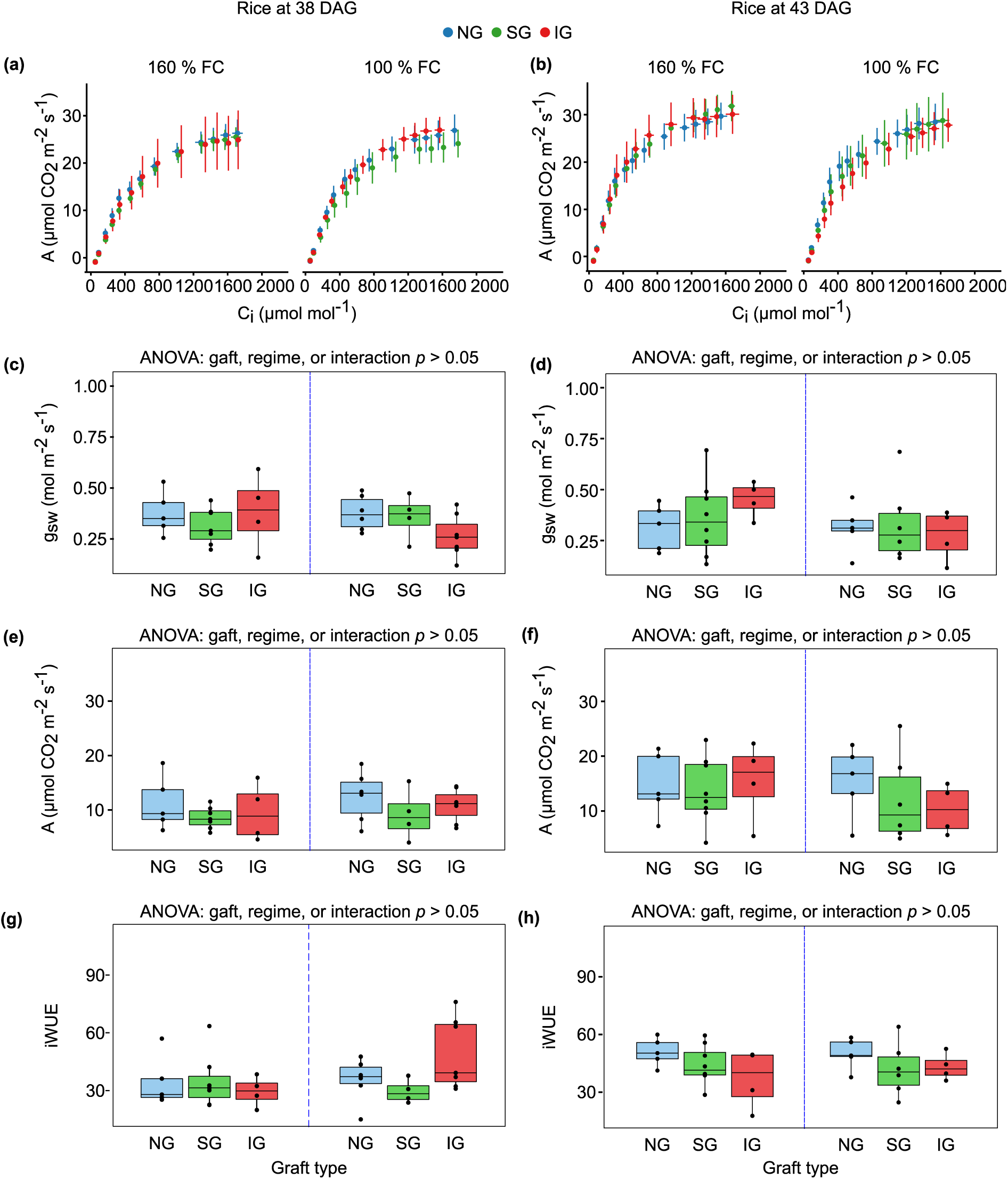
Gas exchange measurements and derived parameters of photosynthetic assimilation in rice plants under contrasting water regimes. CO_2_ response curves were measured in rice non-grafts (NG), self-grafts (SG) and intra-species grafts (IG) at the onset of optimal water regime of 100 % of soil water field capacity (FC) corresponding to 38 days after grafting (DAG) and after five days of sustained optimal water regime, 43 DAG. Control plants under saturated water regime of 160 % FC are shown in the left panels for each respective measurement. Panels **(a, b)** shows net photosynthetic assimilation (*A*) as a function of intercellular CO_2_ (*Ci*) *A/Ci* curve in rice (50 - 2000 µmol mol^−1^). Panels **(c, d)** shows changes in stomatal conductance (*g_sw_*) at 400 µmol mol^−1^. Panels **(e, f)** shows changes in *A* at 400 µmol mol^−1^; and panels **(g, h)** shows intrinsic water use efficiency (*iWUE*) at 400 µmol mol^−1^. Blue, green, and red colors represent NG, SG and IG, respectively. Data in panels **c**-**h** are presented as mean ± SEM (*n*= 4 - 8). Two-way ANOVA was used to evaluate the main effects of graft type and water regime, and their interaction. Where ANOVA results were significant, we detected the statistical differences using Tukey’s HSD post hoc test.

**Fig. 5.**
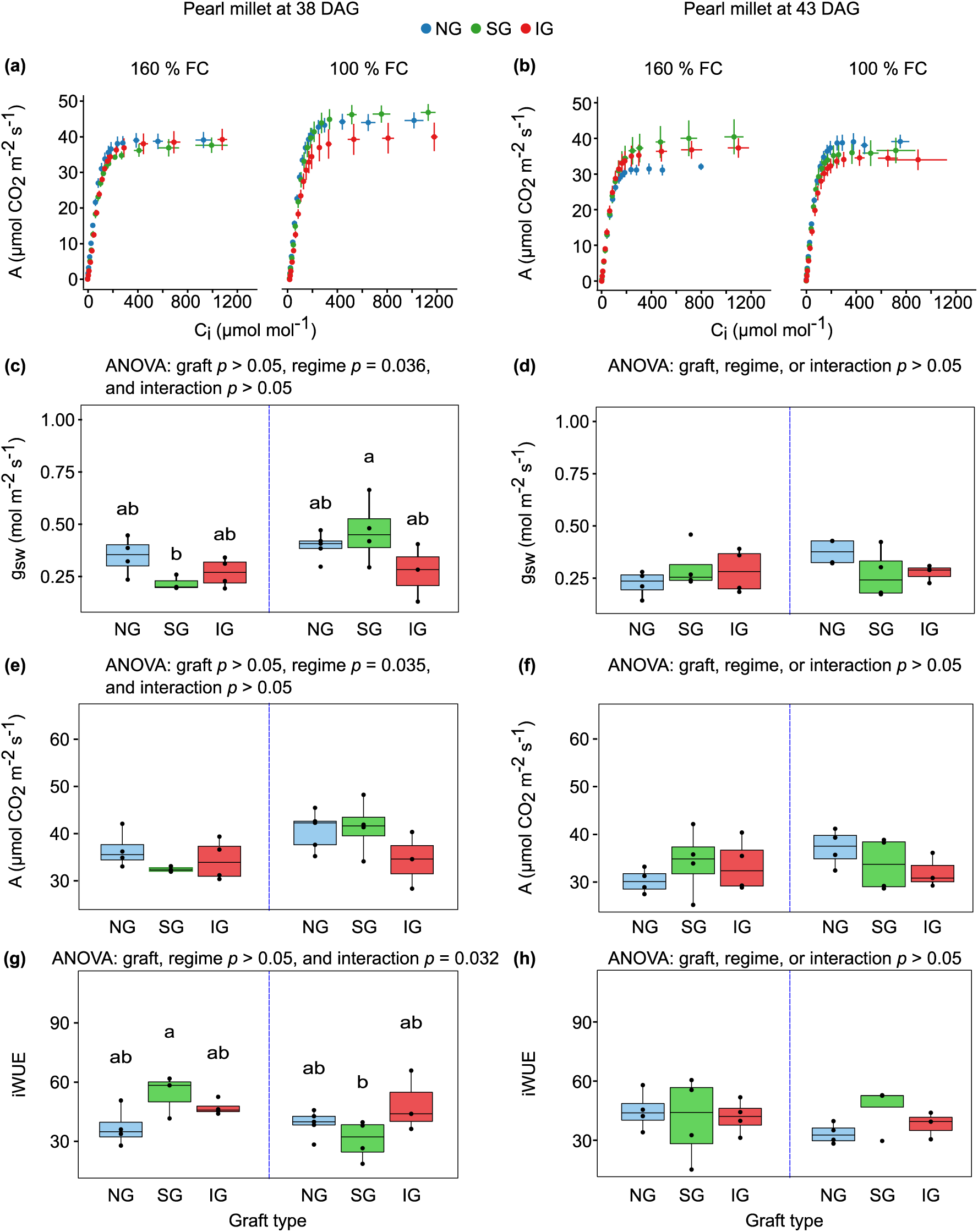
Gas exchange measurements and derived parameters of photosynthetic assimilation in pearl millet plants under contrasting water regimes. CO_2_ response curves were measured in pearl millet non-grafts (NG), self-grafts (SG) and intra-species grafts (IG) at the onset of optimal water regime of 100 % of soil field capacity (FC) corresponding to 38 days after grafting (DAG) and after five days of sustained optimal water regime, 43 DAG. Control plants under saturated water regime of 160 % FC are shown in the left panel for each respective measurement. Panel **(a, b)** shows net photosynthetic assimilation (*A*) as a function of intercellular CO_2_ (*Ci*) *A/Ci* curve in rice (0 - 1250 µmol mol^−1^). Panel **(c, d)** shows changes in stomatal conductance (*g_sw_*) at 400 µmol mol^−1^. Panel **(e, f)** shows changes in *A* at 400 µmol mol^−1^; and panels **(g, h)** shows intrinsic water use efficiency (*iWUE*) at 400 µmol mol^−1^. Blue, green and red colors represent NG, SG and IG, respectively. Data in panels **c** - **h** are presented as mean ± SEM (*n*= 3 - 4). Two-way ANOVA was used to evaluate the main effects of graft type and water regime, and their interaction. Where ANOVA results were significant, we detected the statistical differences using Tukey’s HSD post hoc test.

**Table 2.**
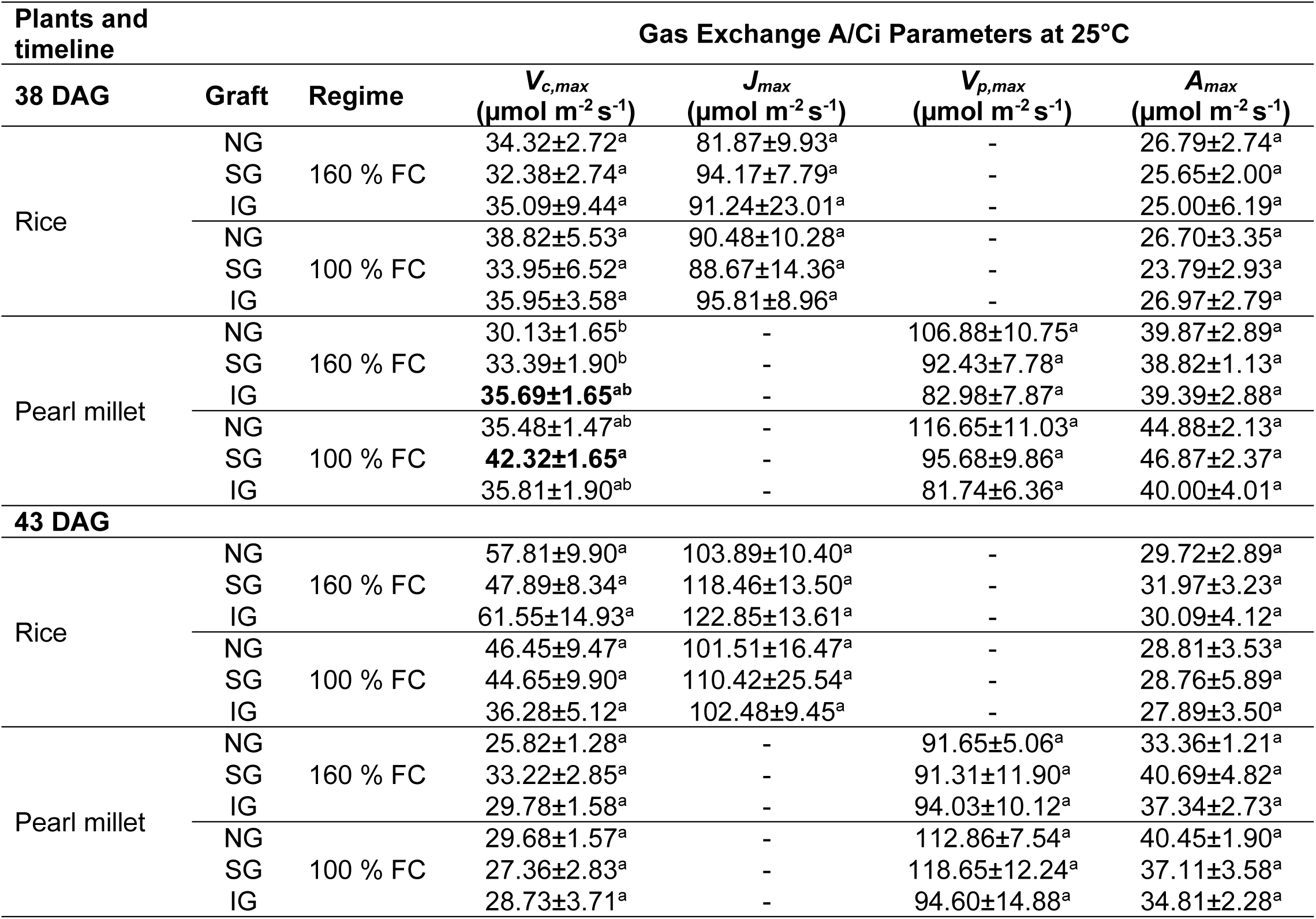
Key photosynthetic parameters of rice and pearl millet under optimal water regime. Key photosynthesis parameters, including maximum carboxylation rate (*V_c,max_*), maximum assimilation rate (*A_max_*), maximum rate of electron transport (*J_max_*) and maximum PEP Carboxylase carboxylation rate (*V_p,max_*) were derived from CO_2_ response curves generated at 25 °C in rice and pearl millet, non-grafts (NG), self-grafts (SG) and intra-species grafts (IG), under saturated water regime of 160 % of soil water field capacity (FC) and at optimal water regime of 100 % FC at 38 and 43 days after grafting (DAG). We derive estimates of the maximum Rubisco carboxylation rate (*V_c,max_*) and maximum assimilation rate (*A_max_*) in both plants. While maximum rate of electron transport (*J_max_*) was estimated in rice only, and the maximum PEP Carboxylase carboxylation rate (*V_p,max_*) in pearl millet. Data are means ± SEM (*n* = 3 - 8). Two-way ANOVA was used to evaluate the effect of graft type on the photosynthesis parameter. Different lowercase superscript letters within columns indicate significant differences (*p* < 0.05) between the graft types as determined by Tukey’s HSD post-hoc test.

### Light-induced stomatal kinetics and dynamic stomatal-photosynthetic coordination are preserved across sustained optimal water regime across graft combinations

To determine whether cereal grafting influences dynamic stomatal behavior during plant development, we examined stomatal responses to step changes in light intensity and quality in rice and pearl millet. We further evaluated whether grafting altered stomatal-photosynthetic coordination across contrasting water regimes, developmental stages and graft combinations (Fig. **6**). Consistent with the observations at 35 DAG, we observed marginal variations in *g_sw_* in both rice and pearl millet plants at 38 and 43 DAG across all graft combinations (Fig. **6a, b**; Table. **S10, S11**). As at 35 DAG, we quantified the dynamic coordination between *g_sw_* and *A* by calculating time constants (τ) required to reach 63 % of the steady-state maximum for both parameters following step changes in light intensity and quality. This analysis allowed us to assess whether grafting altered stomatal and photosynthetic induction kinetics during plant development at 38 DAG (Fig. **6c, d**), and 43 DAG (Fig. **6g, h**). Similar trends were observed at 38 and 43 DAG across all rice graft types. In contrast, pearl millet showed a significant main effect of grafting on *τ_A_* under high red light at 38 DAG (*p* = 0.020), indicating a genotype dependent influence of grafting on photosynthetic induction kinetics during onset of optimal water regime (Table. **S12;** see pearl millet, red light).

**Fig 6.**
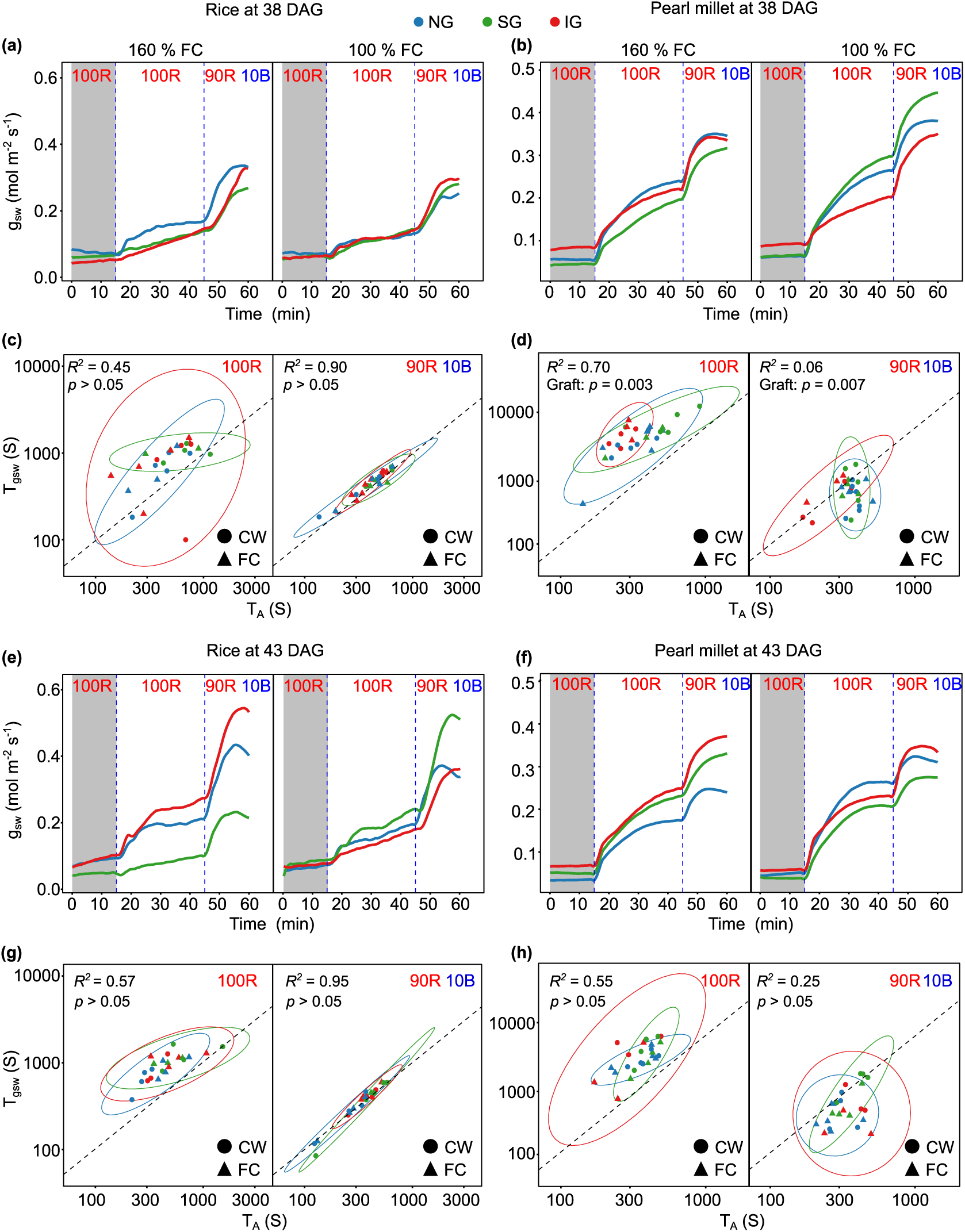
Dynamic photosynthetic and stomatal responses to light induction in rice and pearl millet under contrasting water regimes. The effects of light intensity and quality on stomatal kinetics were assessed in rice and pearl millet, including non-grafts (NG), self-grafts (SG) and intra-species grafts (IG) at the onset of optimal water regime of 100 % of soil water field capacity (FC) corresponding to 38 days after grafting (DAG) and after five days of sustained optimal water regime, 43 DAG. Control plants under the saturated water regime of 160 % FC are shown in the left panels for each respective measurement. Panels **(a**,**b,e,f)** show stomatal conductance (*g_sw_*) response under sequential step-changes in light intensity and quality in rice and pearl millet, respectively. The regions in grey color represent periods of light transition from low light (100 µmol m^−2^ s^−1^) to high light (1500 µmol m^−2^ s^−1^). Variation of light quality was as follows: 100 µmol m^−2^ s^−1^ red light for 15 min, to 1500 µmol m^−2^ s^−1^ red light for 30 min and 1500 µmol m^−2^ s^−1^ and 90:10 red:blue light ratio for 15 min. Panels **(c,d,g,h)** are modelled correlations between the time constants of stomatal conductance (τ_gsw_) and net photosynthetic CO_2_ assimilation (τA). All panels on left show responses following the transition to high red light (1500 µmol m^−2^ s^−1^); while the panels on right show responses following the subsequent transition to 1500 µmol m^−2^ s^−1^ at a 90:10 red:blue ratio. Time constants represent the time required to reach 63 % of the steady-state maximum for each parameter following the light transitions. The dashed line represents the 1:1 coordination line between τ_gsw_ and τA. In the panels the blue, green and red lines represent NG, SG and IG, respectively. Data in panels **a,b,e,f** is presented as mean ± SEM (*n* = 3 - 6).

Following the addition of blue light, stomatal and photosynthetic responses increased in both grafted and non-grafted plants. Grafted plants exhibited statistically similar responses to blue light as non-grafted plants regardless of the species, with few exceptions in pearl millet (Table. **S10-12**; see pearl millet, 43 DAG, blue light). In pearl millet grafting significantly affected *τ_A_* at 38 DAG (main effect of grafting, *p* < 0.001), while *τ_gsw_* at 43 DAG was influenced by both grafting (*p* = 0.009) and water regime (*p* = 0.016) but upon sustained optimal water regime (Table. **S12**; see pearl millet, 43 DAG, graft and regime under blue light). In addition to these kinetic responses, our coordination analyses showed that addition of 10 % blue light substantially reduced the lag between *τ_gsw_* and *τ_A_* that was evident under high red light, indicating improved coordination between stomatal and photosynthetic responses (Fig. **6c, d, g, h**; see all panels of the right side; Table. **S13**). Overall, our results demonstrate that cereal grafting does not disrupt the coupling between light dependent stomatal signaling and photosynthetic induction. Rather, the species-specific patterns of stomatal-photosynthetic coordination observed in rice and pearl millet were maintained across graft combinations, water regimes and plant developmental stages. Although occasional differences in individual kinetic parameters were detected, cereal grafting did not fundamentally alter the stomatal-photosynthetic coordination in either species.

## Discussion

### Graft-induced growth and physiological responses are species-specific

In this study, our experimental design allowed us to investigate whether cereal grafting combined with controlled water regimes, alters plant growth and both steady-state and dynamic physiological performance in rice and pearl millet. Notably, cereal grafting influenced the growth of pearl millet grafts at early stages (Fig. **S1**, Table. **S1, S2**; see 35 and 38 DAG, height), but these effects diminished at later developmental stages (Fig. **1c**; Table **S2**; see 43 DAG, height). Previous studies in dicotyledonous species, for instance *Vitis vinifera*, have shown that grafting same species influence growth (Tedesco *et al*., 2020; Villa-Llop *et al*., 2026), with proposed mechanisms including cambium alignment (Villa-Llop *et al*., 2026) and rootstock-scion interaction (Tedesco *et al*., 2020). Our study extends this body of evidence to cereal crops: although non-significant, rice grafts showed increased growth under controlled water regime at earlier stages (Fig. **S1**), possibly reflecting improved alignment of vascular tissues at the graft junction. This vascular alignment hypothesis is consistent with our finding that, within grafted pearl millet, self-grafted individuals exhibited closer values to non-grafts than intra-species grafts, although these differences were not statistically significant. Although self- and intra-species grafts share the same genetic background, this difference may instead reflect the physical compatibility of excised surfaces. In self-grafts, scion and rootstock originate from complementary halves of the same seed, and their excised surfaces are therefore more likely to match in shape and size, facilitating precise tissue apposition, graft union formation and vascular reconnection. In intra-species grafts, scion and rootstock are derived from two separate seeds, where variation in excised surface and misalignment of embryonic tissues may result in less consistent tissue apposition, contributing to the greater variability observed in this graft-type.

Beyond growth, cereal grafting was associated with physiological adjustments that differed between the two species. In rice, photosynthetic rates increased overtime regardless of water regime, pointing to a robust capacity to maintain carbon assimilation and photosynthetic performance following grafting. In contrast, pearl millet exhibited adjustments in photosynthetic performance in grafted plants during early plant growth stages (35 DAG) up to the onset of optimal water conditions (38 DAG). These changes were accompanied by a simultaneous increase in *iWUE*, suggesting a shift toward resource conservation during this adjustment period. Collectively, our results show that cereal grafting results in species-specific physiological trajectories. We next discuss these physiological adjustments as underpinning steady-state photosynthetic performance and dynamic light-induced stomatal kinetics in rice and pearl millet.

### Species-specific photosynthetic trajectories and transient graft-induced delay in pearl millet

Our results show that grafting has no substantial effect on steady-state photosynthetic performance in rice, contrasting with transient adjustments in stomatal conductance and carbon assimilation observed in pearl millet. This species-specific divergence in grafting outcomes is consistent with the known differences in stomatal regulatory flexibility and photosynthetic biochemistry between C_3_ and C_4_ plants, in which distinct carbon-concentrating mechanisms confer fundamentally different capacities for stomatal and assimilatory regulation (Cui, 2021). At 35 DAG, rice grafts displayed no detectable difference in CO_2_ response (*A/Ci*) curves, photosynthetic biochemical parameter (*V_c.max_, J_max_, A_max_*) or gas exchange traits compared to non-grafts (Fig. **2**). This indicates that cereal grafting does not impair the assimilation, biochemical and CO_2_ diffusional processes, underpinning C_3_ photosynthesis, consistent with observations in dicotyledonous grafts, for example in *V. vinifera* where photosynthetic performance is maintained in well-watered conditions (Opoku *et al*., 2024; Villa-Llop *et al*., 2026). In pearl millet, we observed reduced *A_max_*, *g_sw_* and increased *iWUE* in intra-species grafted plants compared to non-grafts at this early timepoint. These adjustments suggest that cereal grafting transiently modulates stomatal-biochemical coupling in C_4_ species, consistent with the known sensitivity of C_4_ photosynthesis to changes in CO_2_ availability and stomatal regulation (Opoku *et al*., 2024).

When assessed over time corresponding to different plant growth stages, and under contrasting water regimes, these initial differences were no longer apparent, with no consistent differences detected between grafts and non-grafts (Fig. **4,5**; see 43 DAG). Across the study period (35 - 43 DAG), neither rice nor pearl millet showed persistent graft-induced effects on the *A/Ci* response, and the derived key biochemical parameters remained largely conserved between grafted and non-grafted plants. In rice, *V_c.max_*, *J_max_* and *A_max_* increased over time under both water regimes, consistent with overall plant growth, leaf maturation and developmental acclimation (Villa-Llop *et al*., 2026). In contrast, the shifting trajectory of photosynthetic performance in grafted pearl millet over time is consistent with the vascular alignment hypothesis discussed above. Despite the contrasting water regimes used in this study (saturated- and optimal), they were non-limiting for pearl millet, as grafted plants exhibited reduced photosynthetic performance at 35 DAG relative to non-grafted controls, a difference that was no longer apparent by 38 DAG (Fig**. 5**; see 38 DAG). By 43 DAG, self- and intra-species grafts maintained higher but non-significant photosynthetic rates than non-grafts under saturated regime, whose values subsequently followed a typical developmental decline. This pattern suggests that cereal grafting transiently delayed, rather than impaired, the developmental trajectory of photosynthetic capacity in pearl millet, with grafted plants subsequently aligning with and even moderately exceeding non-grafted controls. One possible explanation is that this transient reduction reflects a developmental delay associated with the establishment of functional vascular connections between the grafted embryonic tissues. Unlike dicotyledonous grafting system, cereal grafting relies on *de novo* formation of vascular connections rather than the reconnection of pre-existing mature vasculature (Reeves *et al*., 2022). Consequently, root-to-shoot transport of water, nutrients and signaling molecules may be temporarily constrained until these connections become fully functional, thereby limiting photosynthetic performance during early establishment (Tedesco *et al*., 2020; Camboué *et al*., 2025; Villa-Llop *et al*., 2026). However, this interpretation remains speculative and would require direct assessment of vascular development and hydraulic function, for example through measurements of hydraulic conductance or vascular continuity, to establish a causal link. The absence of lasting graft-induced effects on photosynthetic capacity across both species suggests that cereal grafting system is physiologically robust, whilst the transient nature and species-specificity of early adjustments point to an interaction between graft establishment and the distinct regulatory frameworks of C_3_ and C_4_ photosynthesis.

### Preservation of light quality responses and stomatal-photosynthetic coordination

Rice and pearl millet exhibit distinct stomatal and photosynthetic induction behavior under changing light, and these species-specific patterns remain preserved upon cereal grafting and across water regimes. Dynamic stomatal responses to step changes in light intensity and quality support this conclusion, as they allowed separation of the photosynthesis-driven and phototropin-mediated components of stomatal opening (Matthews *et al*., 2020). Across all timepoints and water regimes, all graft types of rice and pearl millet showed slow stomatal and photosynthetic induction under red light, followed by marked acceleration under blue light (Fig. **3**,**6**, Tables **S5-6, S10-11**). This lag is consistent with established guard-cell signaling pathways, in which red light triggers stomatal opening indirectly via photosynthesis-dependent reductions in intercellular CO_2_ (Vialet-Chabrand *et al*., 2021), while blue light activates a guard cell signaling cascade mediated by phototropins and the plasma membrane *H^+^-ATPase* (Matthews *et al*., 2020). Blue light triggered distinct stomatal-photosynthetic coordination strategies between the two species. Under the red:blue light, the speed of stomatal opening (τ_gsw_) and photosynthetic induction (τA) converged toward the 1:1 coordination line across all rice graft types, indicating tightly synchronized activation of the Calvin-Benson Cycle and phototropin-mediated stomatal opening (Matthews *et al*., 2020). In contrast, pearl millet graft types consistently exhibited τ_gsw_ values lower than τA, indicating that the rate-limiting step during light transitions is biochemical activation rather than diffusional constraints imposed by stomata, a decoupling characteristic of C_4_ physiology, where carbon-concentrating mechanisms reduce dependence on stomatal aperture to maintain internal CO_2_ supply (Zhen & Bugbee, 2020).

Cereal grafting did not alter induction kinetics or the signaling architecture linking light perception to gas exchange responses in either species. Although pearl millet grafts showed minor, transiently slower induction at 38 DAG, species-specific stomatal-photosynthetic coordination was maintained regardless of graft type across all measured timepoints and water regimes. This confirms that graft union formation and vascular connection inherent to cereal grafting does not disrupt critical guard-cell signaling pathways, including phototropin-mediated blue-light responses and red light-driven photosynthetic coupling.

Taken together, these results indicate that cereal grafting does not fundamentally alter the species-specific physiological strategies governing stomatal-photosynthetic coordination. Rice physiological traits appear largely insensitive to the cereal grafting, while pearl millet undergoes a brief, self-resolving adjustment period that does not compromise its underlying biochemical capacity or long-term physiological coordination. Whether this differential sensitivity reflects differences in vascular anatomy, graft union dynamics or developmental timing between C_3_ and C_4_ cereals remains an open question for future investigation.

## Conclusion

Our findings establish cereal grafting as a physiologically robust platform for investigating root-shoot communication in cereals. By demonstrating that grafting itself does not substantially alter the traits most closely linked to carbon gain and water-use, our work provides an essential foundation for future studies aimed at dissecting long-distance signaling networks, hydraulic regulation and hormonal control of crop performance. More broadly, the ability to combine genetically distinct root and shoot systems without compromising physiological function creates new opportunities to explore trait interactions, exploit natural variation and accelerate the development of climate-resilient cereals. As grafting technologies become increasingly tractable in monocots, this approach offers a powerful experimental and translational framework for understanding and engineering complex traits in global crop staples.

## Acknowledgements

The work was supported by UKRI-Future Leaders Fellowship MR/Y016882/1; *REVOLUTION* to Pallavi Singh. Anoop Tripathi was supported by a Gates Cambridge Trust PhD Student Fellowship. We acknowledge Mouesanao K. Kandjoze and Urvi K. Gandhi for their assistance with plant growth, plant maintenance, and physiological measurements.

## Authors contributions

**PS** conceptualized the project, obtained the funding and supervised the work. **CMM** designed the study. **CMM** and **XSMG**, carried out the experimental work, optimized data analysis pipelines, analyzed the data and interpreted the results. **AT** grafted plants. **XSMG** conducted physiological and gas exchange measurements. **CMM**, **XSMG** and **PS** prepared the figures and wrote the article. **CMM, XSGM, AT and PS** contributed to the final version of the manuscript and approved it for publication.

## Data availability

All data needed to evaluate the conclusions in the paper are present in the paper and / or the Supporting Information.

