## Supporting Information for "Cereal grafting in rice and pearl millet preserves photosynthetic performance and stomatal dynamics, establishing a platform for root-shoot communication studies"

Article acceptance date: Click here to enter a date.

The following Supporting Information is available for this article:

**Fig S1.** Plant growth measurements of rice and pearl millet plants at 35 and 38 days after grafting

**Table S1.** Two-way ANOVA results on rice plants growth traits: biomass and height.

**Table S2.** Two-way ANOVA results on pearl millet plants growth traits: biomass and height

**Table S3.** One-way ANOVA results on photosynthesis biochemical parameters of rice and pearl millet.

**Table S4.** One-way ANOVA results on core parameters of gas exchange in rice and pearl millet

**Table S5.** Time constants for stomatal conductance ($\tau_{gsw}$) and net $CO_2$ assimilation rate ($\tau_A$) during step changes in light intensity and quality in rice and pearl millet

**Table S6.** Time constants for stomatal conductance ($\tau_{gsw}$) and net $CO_2$ assimilation rate ($\tau_A$) during step changes in light intensity and quality in rice and pearl millet

**Table S7.** One-way ANOVA results on the photosynthetic-stomatal coordination index ($\tau_{gsw}/\tau_A$) during step changes in light intensity and quality in rice and pearl millet.

**Table S8.** Two-way ANOVA results on photosynthesis biochemical parameters of rice and pearl millet.

**Table S9.** Two-way ANOVA results on core parameters of gas exchange in rice and pearl millet.

**Table S10.** Time constants for stomatal conductance ($\tau_{gsw}$) and net $CO_2$ assimilation rate ($\tau_A$) during step changes in light intensity and quality in rice.

**Table S11.** Time constants for stomatal conductance ($\tau_{gsw}$) and net $CO_2$ assimilation rate ($\tau_A$) during step changes in light intensity and quality in pearl millet.

**Table S12.** Two-way ANOVA results on time constants for stomatal conductance ($\tau_{gsw}$) and net $CO_2$ assimilation rate ($\tau_A$) during step changes in light intensity and quality in rice and pearl millet.

**Table S13.** Two-way ANOVA results on the photosynthetic-stomatal coordination index ($\tau_{gsw}/\tau_A$) during step changes in light intensity and quality in rice and pearl millet.

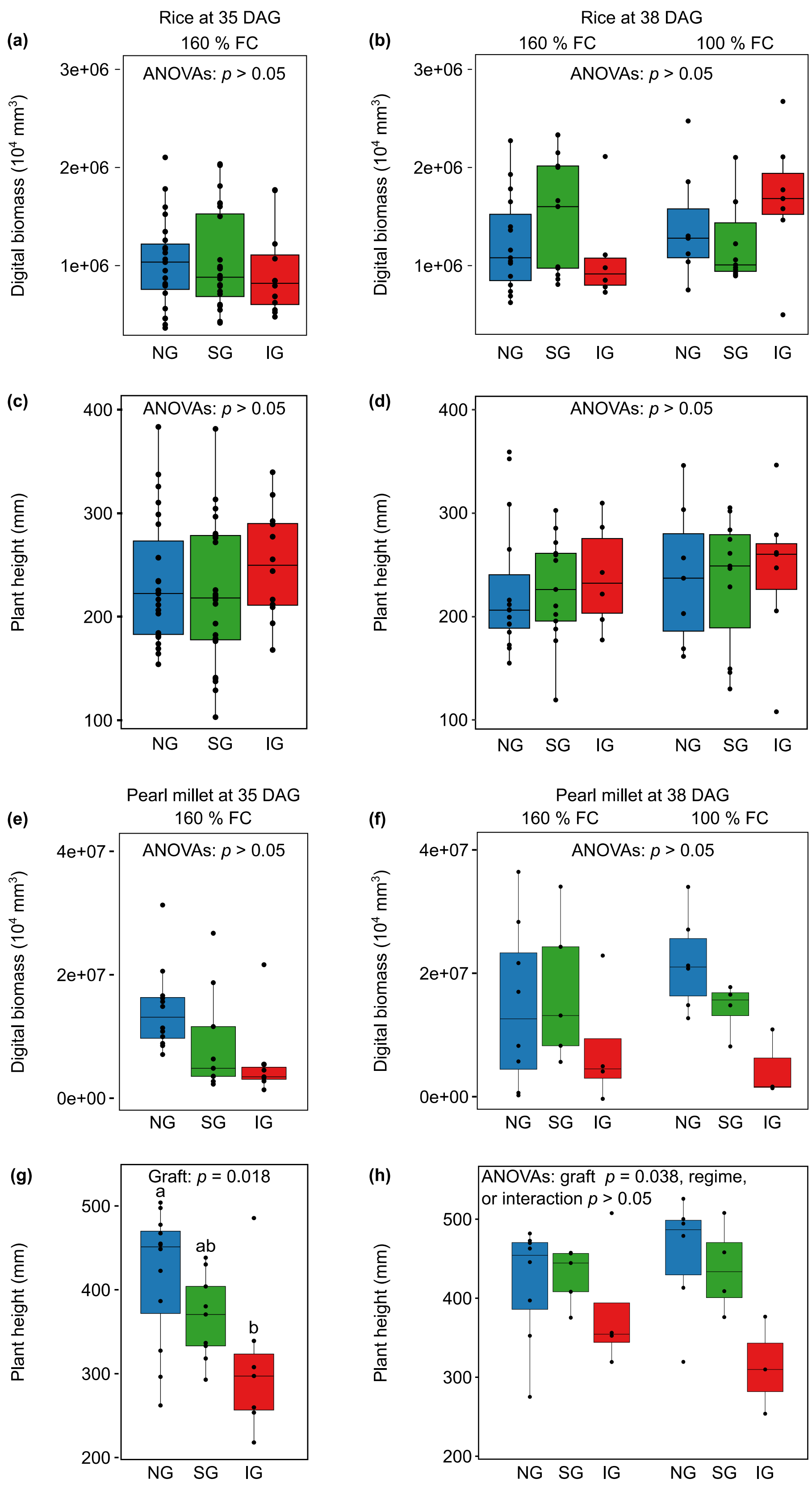
Rice at 35 DAG
160 % FC
Rice at 38 DAG
160 % FC
100 % FC
(a)
(b)
(c)
(d)
ANOVAs: p > 0.05
Digital biomass (10⁴ mm³)
Plant height (mm)
NG
SG
IG
Pearl millet at 35 DAG
160 % FC
Pearl millet at 38 DAG
160 % FC
100 % FC
(e)
(f)
(g)
(h)
Graft: p = 0.018
a
ab
b
ANOVAs: graft p = 0.038, regime,
or interaction p > 0.05

**Fig. S1. Plant growth measurements of rice and pearl millet plants at 35 and 38 days after grafting**. Plant digital biomass and height were measured on rice and pearl millet non-grafts (NG), self-graft (SG) and Intra-species graft (IG), grown at saturated water regime (160 % of soil water field capacity) at 35 days after grafting (DAG), and at onset of optimal water regime 100 %FC, 38 DAG. Panels **a, c** and **b,d** are rice plants at 35 and 38 DAG, respectively. Panels **e,g** and **f,h** are pearl millet plants at 35, and 38 DAG respectively. Data is presented as mean ± SEM (*n* = 3-16). Two-way ANOVA was used to evaluate the main effect of graft type and water regime, and their interactions.

**Table S1. Two-way ANOVA results on rice growth traits: biomass and height.** Digital images were used to estimate biomass and height of rice plants including non-grafts (NG), self-grafts (SG) and intra-species grafts (IG), under saturated water regime of 160 % of soil water field capacity (FC) and at optimal water regime of 100 % FC across timepoints: 35, 38, and 43 days after grafting. The digital images were captured using PlantEye. Data are the mean ± standard error ($n$ = 6 - 24). Data were analyzed using two-way ANOVA linear model consisting of graft type, water regime and their interaction as model explanatory factors. Difference between the groups were detected by Tukeys HSD test for multiple comparisons at $p \leq 0.05$. Statistically significant effects are indicated in bold. DAG, stands for days after grafting; Df(n,d), stands for the degree of freedom, numerator, and denominator; F stands for F test value; P stands for probability value in at ≤ 0.05.

| Time point | Source of variation | Biomass | | | Height | | |
|---|---|---|---|---|---|---|---|
| | | **Df(n,d)** | **F** | **P** | **Df(n,d)** | **F** | **P** |
| 35 DAG | Graft | 2,56 | 0.234 | 0.792 | 2,56 | 0.822 | 0.444 |
| 38 DAG | Graft | 2,53 | 0.268 | 0.766 | 2,53 | 0.086 | 0.918 |
| | Regime | 1,53 | 0.236 | 0.629 | 1,53 | 0.187 | 0.668 |
| | Graft*Regime | 2,53 | 2.940 | 0.062 | 2,53 | 0.047 | 0.954 |
| 43 DAG | Graft | 2,52 | 0.536 | 0.588 | 2,52 | 0.220 | 0.804 |
| | Regime | 1,52 | 0.303 | 0.584 | 1,52 | 0.769 | 0.385 |
| | Graft*Regime | 2,52 | 1.785 | 0.178 | 2,52 | 0.105 | 0.900 |

**Table S2. Two-way ANOVA results on pearl millet growth traits: biomass and height.** Digital images were used to estimate biomass and height of rice plants including non-grafts (NG), self-grafts (SG) and intra-species grafts (IG), under saturated water regime of 160 % of soil water field capacity (FC) and at optimal water regime of 100 % FC across timepoints: 35, 38, and 43 days after grafting. The digital images were captured using PlantEye. Data are the mean ± standard error (*n* = 3 - 12). Data were analyzed using two-way ANOVA linear model consisting of graft type, water regime and their interaction as model explanatory factors. Difference between the groups were detected by Tukeys HSD test for multiple comparisons at *p* ≤ 0.05. Statistically significant effects are indicated in bold. Df(n,d), stands for the degree of freedom, numerator, and denominator; F stands for F test value; P stands for probability value in at ≤ 0.05.

| Time point | Source of variation | Biomass | | | Height | | |
|---|---|---|---|---|---|---|---|
| | | **Df(n,d)** | **F** | **P** | **Df(n,d)** | **F** | **P** |
| 35 DAG | Graft | 2,25 | 3.050 | 0.065 | 2,25 | 4.706 | **0.018** |
| 38 DAG | Graft | 2,24 | 2.902 | 0.074 | 2,24 | 3.750 | **0.038** |
| | Regime | 1,24 | 0.193 | 0.664 | 1,24 | 0.014 | 0.907 |
| | Graft*Regime | 2,24 | 0.861 | 0.435 | 2,24 | 1.406 | 0.265 |
| 43 DAG | Graft | 2,23 | 1.076 | 0.357 | 2,23 | 2.728 | 0.086 |
| | Regime | 1,23 | 1.062 | 0.314 | 1,23 | 0.114 | 0.739 |
| | Graft*Regime | 2,23 | 0.046 | 0.955 | 2,23 | 0.232 | 0.795 |

**Table S3. One-way ANOVA results on photosynthesis biochemical parameters of rice and pearl millet**. Key photosynthesis parameters, including maximum carboxylation rate (*$V_{c,max}$*), maximum PEP Carboxylase carboxylation rate (*$V_{p,max}$*), maximum electron transport rate (*$J_{max}$*), and maximum net photosynthesis rate (*$A_{max}$*) were derived from $CO_2$ response curves of rice and pearl millet's non-grafts (NG), self-grafts (SG) and intra-species grafts (IG), under saturated water regime of 160 % of soil water field capacity at 35 days after grafting. Data are the mean ± standard error (*n* = 6 - 11). Data were analyzed using one-way ANOVA linear model consisting of graft type as model explanatory factors. Difference between the groups were detected by Tukeys HSD test for multiple comparisons at *p* ≤ 0.05. Statistically significant effects are indicated in bold. DAG, stands for days after grafting; Df(n,d), stands for the degree of freedom, numerator, and denominator; F stands for F test value; P stands for probability value in at ≤ 0.05.

| | | | ***$V_{c,max}$*** | | | ***$J_{max}$*** | | | ***$V_{p,max}$*** | | | ***$A_{max}$*** | | |
|---|---|---|---|---|---|---|---|---|---|---|---|---|---|---|
| **Plant** | **Timepoint** | **Source of variation** | **Df(n,d)** | **F** | **P** | **Df(n,d)** | **F** | **P** | **Df(n,d)** | **F** | **P** | **Df(n,d)** | **F** | **P** |
| Rice | 35 DAG | Graft | 2,27 | 0.656 | 0.527 | 2,23 | 0.116 | 0.891 | - | - | - | 2,27 | 0.640 | 0.535 |
| Pearl millet | | | 2,20 | 0.854 | 0.441 | - | - | - | 2,20 | 2.247 | 0.891 | 2,20 | 5.685 | **0.011** |

**Table S4. One-way ANOVA results on core parameters of gas exchange in rice and pearl millet.** Key gas exchange parameter, including net CO2 assimilation rate (*A*), stomatal conductance to water vapor (*gsw*), and intrinsic water use efficiency (*iWUE*) were derived $CO_2$ response of rice and pearl millet's non-grafts (NG), self-grafts (SG) and intra-species grafts (IG), under saturated water regime of 160 % of soil water field capacity at 35 days after grafting. Data are the mean ± standard error (*n* = 6 - 11). Data were analyzed using one-way ANOVA linear model consisting of graft type as model explanatory factors. Difference between the groups were detected by Tukeys HSD test for multiple comparisons at $p \leq 0.05$. Statistically significant effects are indicated in bold. DAG, stands for days after grafting; Df(n,d), stands for the degree of freedom, numerator, and denominator; F stands for F test value; P stands for probability value in at ≤ 0.05.

| **Plant** | **Timepoint** | **Source of variation** | ***A*** | | | ***$g_{sw}$*** | | | ***iWUE*** | | |
|---|---|---|---|---|---|---|---|---|---|---|---|
| | | | **Df(n,d)** | **F** | **P** | **Df(n,d)** | **F** | **P** | **Df(n,d)** | **F** | **P** |
| Rice | 35 DAG | Graft | 2,27 | 1.396 | 0.265 | 2,27 | 0.714 | 0.499 | 2,27 | 0.745 | 0.484 |
| Pearl millet | | | 2,20 | 4.925 | **0.018** | 2,20 | 5.800 | **0.010** | 2,20 | 4.689 | **0.021** |

**Table S5. Time constants for stomatal conductance ($\tau_{gsw}$) and net $CO_2$ assimilation rate ($\tau_A$) during step changes in light intensity and quality in rice and pearl millet.** We estimated in rice non-grafts (NG), self-grafts (SG) and intra-species grafts (IG), under saturated water regime of 160 % of soil water field capacity at 35 days after grafting, the time constants (the time required to reach 63 % of the steady-state maximum) for stomatal conductance ($g_{sw}$) and net $CO_2$ assimilation ($A$) following step changes in light intensity (red light; from 100 to 1500 μmol $m^{-2}$ $s^{-1}$) and a change in light quality (blue light; from 1500 μmol $m^{-2}$ $s^{-1}$ red light to 1500 μmol $m^{-2}$ $s^{-1}$ composed of 90 % red and 10 % blue light). Data are means ± SEM ($n$ = 5 - 10). Data were analyzed using one-way ANOVA linear model consisting of graft type as model explanatory factors. Difference between the groups were detected by Tukeys HSD test for multiple comparisons at $p \leq 0.05$. Different lowercase superscript letters within columns indicate significant differences between groups as determined by Tukey's HSD post-hoc test. DAG, stands for days after grafting.

| Plants | Light | Timepoint | Graft | $\tau_{gsw}$ (S) | Duration (min) | $\tau_A$ (S) | Duration (min) |
|---|---|---|---|---|---|---|---|
| Rice | 35 DAG | Red light | NG | 1092.96 ± 97.19$^{a}$ | 18 min 12 s | 476.80 ± 97.04$^{a}$ | 7 min 56 s |
| | | | SG | 1072.01 ± 53.64$^{a}$ | 17 min 52 s | 670.18 ± 87.81$^{a}$ | 11 min 10 s |
| | | | IG | 995.72 ± 80.53$^{a}$ | 16 min 35 s | 466.33 ± 59.27$^{a}$ | 7 min 46 s |
| | | Blue light | NG | 504.68 ± 87.95$^{a}$ | 8 min 24 s | 464.12 ± 68.79$^{a}$ | 7 min 44 s |
| | | | SG | 475.30 ± 36.95$^{a}$ | 7 min 55 s | 435.94 ± 24.66$^{a}$ | 7 min 15 s |
| | | | IG | 479.77 ± 24.54$^{a}$ | 7 min 59 s | 505.46 ± 43.63$^{a}$ | 8 min 25 s |
| Pearl millet | | Red light | NG | 708.85 ± 62.11$^{a}$ | 11 min 48 s | 375.24 ± 51.81$^{a}$ | 6 min 15 s |
| | | | SG | 712.28 ± 19.27$^{a}$ | 11 min 52 s | 423.15 ± 25.36$^{a}$ | 7 min 3 s |
| | | | IG | 653.54 ± 31.78$^{a}$ | 10 min 53 s | 317.21 ± 34.28$^{a}$ | 5 min 17 s |
| | | Blue light | NG | **266.06 ± 20.04$^{a}$** | 4 min 26 s | **370.38 ± 16.99$^{a}$** | 6 min 10 s |
| | | | SG | 207.26 ± 10.36$^{b}$ | 3 min 27 s | 294.26 ± 15.74$^{b}$ | 4 min 54 s |
| | | | IG | 226.76 ± 19.16$^{ab}$ | 3 min 46 s | 268.43 ± 19.15$^{b}$ | 4 min 28 s |

**Table S6. One-way ANOVA results on time constants for stomatal conductance ($\tau_{gsw}$) and net CO2 assimilation rate ($\tau_A$) during step changes in light intensity and quality in rice and pearl millet.** We estimated in rice and pearl millet non-grafts (NG), self-grafts (SG) and intra-species grafts (IG), under saturated water regime of 160 % of soil water field capacity at 35 days after grafting, the time constants (the time required to reach 63 % of the steady-state maximum) for stomatal conductance ($g_{sw}$) and net $CO_2$ assimilation ($A$) following step changes in light intensity (red light; from 100 to 1500 µmol $m^{-2}$ $s^{-1}$) and a change in light quality (blue light; from 1500 µmol $m^{-2}$ $s^{-1}$ red light to 1500 µmol $m^{-2}$ $s^{-1}$ composed of 90 % red and 10 % blue light). Data are means ± SEM ($n$ = 5 - 10). Data were analyzed using one-way ANOVA linear model consisting of graft type as model explanatory factors. Statistically significant effects are indicated in bold. DAG, stands for days after grafting; Df(n,d), stands for the degree of freedom, numerator, and denominator; F stands for F test value; P stands for probability value in at ≤ 0.05.

| Plant | Light | Timepoint | Source of variation | $\tau_A$ | | | $\tau_{gsw}$ | | |
|---|---|---|---|---|---|---|---|---|---|
| | | | | Df(n,d) | F | P | Df(n,d) | F | P |
| Rice | Red light | 35 DAG | Graft | 2,16 | 0.612 | 0.055 | 2,16 | 0.302 | 0.743 |
| Pearl millet | | | | 2,22 | 1.158 | 0.333 | 2,22 | 0.301 | 0.743 |
| Rice | Blue light | | | 2,16 | 1.594 | 0.234 | 2,16 | 0.332 | 0.722 |
| Pearl millet | | | | 2,22 | 7.093 | **0.004** | 2,22 | 4.995 | **0.016** |

**Table S7. Two-way ANOVA results on the photosynthetic-stomatal coordination index ($\tau_{gsw}/\tau_A$) during step changes in light intensity and quality in rice and pearl millet**. We calculated the coordination index as the ratio between the time constants for stomatal conductance ($\tau_{gsw}$) and net $CO_2$ assimilation rate ($\tau_A$) in rice and pearl millet's non-grafts (NG), self-grafts (SG) and intra-species grafts (IG) under a saturated water regime of 160 % of soil water field capacity at 35 days after grafting. Time constants represent the time required to reach 63 % of the steady-state maximum following step changes in light intensity (red light; from 100 to 1500 µmol $m^{-2}$ $s^{-1}$) and a change in light quality (blue light; from 1500 µmol $m^{-2}$ $s^{-1}$ red light to 1500 µmol $m^{-2}$ $s^{-1}$ composed of 90 % red and 10 % blue light). Data are means ± SEM ($n$ = 5 - 10). Data were analyzed using a one-way ANOVA linear model consisting of graft type as an explanatory factor. Statistically significant effects $p \leq 0.05$ are indicated in bold. DAG, stands for days after grafting; Df(n,d), stands for the degree of freedom, numerator, and denominator; F stands for F test value; P stands for probability value in at ≤ 0.05.

| Plant | Timepoint | Source of variation | Coordination index | | | | | |
|---|---|---|---|---|---|---|---|---|
| | | | Red light | | | Blue light | | |
| | | | Df(n,d) | F | P | Df(n,d) | F | P |
| Rice | 35 DAG | Graft | 2,16 | 2.308 | 0.132 | 2,16 | 2.720 | 0.096 |
| Pearl millet | | | 2,22 | 3.466 | **0.049** | 2,22 | 2.273 | 0.127 |

**Table S8. Two-way ANOVA results on photosynthesis biochemical parameters of rice and pearl millet**. Key photosynthesis parameters, including maximum carboxylation rate ($V_{c,max}$), maximum PEP Carboxylase carboxylation rate ($V_{p,max}$), maximum electron transport rate ($J_{max}$), and maximum net photosynthesis rate ($A_{max}$) were derived from $CO_2$ response curves of rice and pearl millet's non-grafts (NG), self-grafts (SG) and intra-species grafts (IG), under saturated water regime of 160 % of soil water field capacity (FC) and at optimal water regime of 100 % FC at 38 and 43 days after grafting. Data are the mean ± standard error ($n$ = 3 - 8). Data were analyzed using two-way ANOVA linear model with graft type, water regime and their interactions as the main explanatory factors. Statistically significant effects are indicated in bold. DAG, stands for days after grafting; Df(n,d), stands for the degree of freedom, numerator, and denominator; F stands for F test value; P stands for probability value in at ≤ 0.05.

| Plant | Timepoint | Source of variation | $V_{c,max}$ | | | $J_{max}$ | | | $V_{p,max}$ | | | $A_{max}$ | | |
|---|---|---|---|---|---|---|---|---|---|---|---|---|---|---|
| | | | Df(n,d) | F | P | Df(n,d) | F | P | Df(n,d) | F | P | Df(n,d) | F | P |
| Rice | 38 DAG | Graft | 2,27 | 0.328 | 0.723 | 2,22 | 0.144 | 0.867 | | | | 2,27 | 0.159 | 0.853 |
| | | Regime | 1,27 | 0.336 | 0.567 | 1,22 | 0.065 | 0.801 | | | | 1,27 | 4.343 | 0.998 |
| | | Graft*Regime | 2,27 | 0.076 | 0.927 | 2,22 | 0.150 | 0.861 | | | | 2.27 | 0.163 | 0.850 |
| | 43 DAG | Graft | 2,26 | 0.177 | 0.839 | 2,18 | 0.349 | 0.710 | | | | 2,26 | 0.092 | 0.912 |
| | | Regime | 1,26 | 1.959 | 0.173 | 1,18 | 0.471 | 0.501 | | | | 1,26 | 0.427 | 0.519 |
| | | Graft*Regime | 2,26 | 0.587 | 0.563 | 2,18 | 0.142 | 0.868 | | | | 2,26 | 0.041 | 0.960 |
| Pearl millet | 38 DAG | Graft | 2,17 | 5.286 | **0.016** | | | | 2,17 | 4.870 | **0.021** | 2,17 | 1.192 | 0.328 |
| | | Regime | 1,17 | 12.24 | **0.003** | | | | 1,17 | 0.300 | 0.591 | 1,17 | 5.023 | **0.039** |
| | | Graft*Regime | 2,17 | 3.101 | 0.071 | | | | 2,17 | 0.162 | 0.852 | 2,17 | 1.011 | 0.385 |
| | 43 DAG | Graft | 2,17 | 0.628 | 0.546 | | | | 2,17 | 0.549 | 0.587 | 2,17 | 0.404 | 0.674 |
| | | Regime | 1,17 | 0.284 | 0.601 | | | | 1,17 | 4.062 | 0.059 | 1,17 | 0.035 | 0.854 |
| | | Graft*Regime | 2,17 | 2.259 | 0.135 | | | | 2,17 | 0.859 | 0.441 | 2,17 | 1.886 | 0.182 |

**Table S9. Two-way ANOVA results on core parameters of gas exchange in rice and pearl millet.** Key gas exchange parameter, including net $CO_2$ assimilation rate (*A*), stomatal conductance to water vapor ($g_{sw}$), and intrinsic water use efficiency (*iWUE*) were derived $CO_2$ response of rice and pearl millet's non-grafts (NG), self-grafts (SG) and intra-species grafts (IG), under saturated water regime of 160 % of soil water field capacity (FC) and at optimal water regime of 100 % FC at 38 and 43 days after grafting. Data are the mean ± standard error (*n* = 3 - 4). Data were analyzed using two-way ANOVA linear model with graft type, water regime and their interactions as the main explanatory factors. Statistically significant effects are indicated in bold. DAG, stands for days after grafting; Df(n,d), stands for the degree of freedom, numerator, and denominator; F stands for F test value; P stands for probability value in at ≤ 0.05.

| Plant | Timepoint | Source of variation | *A* | | | $g_{sw}$ | | | *iWUE* | | |
|---|---|---|---|---|---|---|---|---|---|---|---|
| | | | Df(n,d) | F | P | Df(n,d) | F | P | Df(n,d) | F | P |
| Rice | 38 DAG | Graft | 2,27 | 0.978 | 0.389 | 2,27 | 1.106 | 0.345 | 2,27 | 1.338 | 0.280 |
| | | Regime | 1,27 | 0.129 | 0.722 | 1,27 | 0.359 | 0.554 | 1,27 | 1.057 | 0.313 |
| | | Graft*Regime | 2,27 | 0.007 | 0.993 | 2,27 | 1.528 | 0.235 | 2,27 | 2.385 | 0.111 |
| | 43 DAG | Graft | 2,26 | 0.144 | 0.867 | 2,26 | 0.266 | 0.769 | 2,26 | 2.301 | 0.120 |
| | | Regime | 1,26 | 0.632 | 0.434 | 1,26 | 1.095 | 0.305 | 1,26 | 0.008 | 0.930 |
| | | Graft*Regime | 2,26 | 0.374 | 0.692 | 2,26 | 0.842 | 0.442 | 2,26 | 0.425 | 0.659 |
| Pearl millet | 38 DAG | Graft | 2,17 | 1.964 | 0.171 | 2,17 | 2.402 | 0.121 | 2,17 | 2.132 | 0.149 |
| | | Regime | 1,17 | 5.257 | **0.035** | 1,17 | 5.185 | **0.036** | 1,17 | 2.136 | 0.162 |
| | | Graft*Regime | 2,17 | 1.693 | 0.213 | 2,17 | 2.974 | 0.078 | 2,17 | 4.242 | **0.032** |
| | 43 DAG | Graft | 2,17 | 0.097 | 0.908 | 2,17 | 0.099 | 0.906 | 2,17 | 0.357 | 0.705 |
| | | Regime | 1,17 | 0.763 | 0.394 | 1,17 | 1.132 | 0.302 | 1,17 | 0.389 | 0.541 |
| | | Graft*Regime | 2,17 | 1.647 | 0.222 | 2,17 | 2.474 | 0.114 | 2,17 | 1.095 | 0.357 |

**Table S10. Time constants for stomatal conductance ($τ_{gsw}$) and net $CO_2$ assimilation rate ($τ_A$) during step changes in light intensity and quality in rice.** We estimated in rice non-grafts (NG), self-grafts (SG) and intra-species grafts (IG), under saturated water regime of 160 % of soil water field capacity (FC) and at optimal water regime of 100 % FC at 38 and 43 days after grafting, the time constants (the time required to reach 63 % of the steady-state maximum) for stomatal conductance ($g_{sw}$) and net $CO_2$ assimilation ($A$) following step changes in light intensity (red light; from 100 to 1500 μmol $m^{-2}$ $s^{-1}$) and a change in light quality (blue light; from 1500 μmol $m^{-2}$ $s^{-1}$ red light to 1500 μmol $m^{-2}$ $s^{-1}$ composed of 90 % red and 10 % blue light). Data are means ± SEM ($n$ = 3 - 6). Data were analyzed using two-way ANOVA linear model with graft type, water regime and their interactions as the main explanatory factors. Difference between the groups were detected by Tukeys HSD test for multiple comparisons at $p \leq 0.05$. Different lowercase superscript letters within columns indicate significant differences ($p \leq 0.05$) between graft as determined by Tukey's HSD post-hoc test. DAG, stands for days after grafting.

| Timepoint | Light | Regime | Graft | $τ_{gsw}$ (S) | Duration (min) | $τ_A$ (S) | Duration (min) |
|---|---|---|---|---|---|---|---|
| 38 DAG | Red light | Saturated water regime (160 % FC) | NG | 706.14 ± 151.34[a] | 11 min 46 s | 452.04 ± 87.07[a] | 7 min 32 s |
| | | | SG | 1024.38 ± 108.38[a] | 17 min 4 s | 736.6 ± 152.09[a] | 12 min 16 s |
| | | | IG | 856.47 ± 269.9[a] | 14 min 16 s | 610.26 ± 85.45[a] | 10 min 10 s |
| | Blue light | | NG | 403.85 ± 64.94[a] | 6 min 43 s | 376.52 ± 69.9[a] | 6 min 16 s |
| | | | SG | 540.72 ± 72.5[a] | 9 min 0 s | 507.01 ± 76.38[a] | 8 min 27 s |
| | | | IG | 588.76 ± 16.5[a] | 9 min 48 s | 537.32 ± 12.52[a] | 8 min 57 s |
| | Red light | Optimal water regime (100% FC) | NG | 688.92 ± 226.99[a] | 11 min 28 s | 533.07 ± 170.09[a] | 8 min 53 s |
| | | | SG | 1052.67 ± 43.35[a] | 17 min 32 s | 558.3 ± 179.01[a] | 9 min 18 s |
| | | | IG | 881.55 ± 199.26[a] | 14 min 41 s | 441.08 ± 105.86[a] | 7 min 21 s |
| | Blue light | | NG | 460.77 ± 99.12[a] | 7 min 40 s | 441.46 ± 95.41[a] | 7 min 21 s |
| | | | SG | 451.27 ± 20.59[a] | 7 min 31 s | 481.89 ± 53.11[a] | 8 min 1 s |
| | | | IG | 413.69 ± 49.64[a] | 6 min 53 s | 378.02 ± 44.42[a] | 6 min 18 s |
| 43 DAG | Red light | Saturated water regime (160 % FC) | NG | 581.51 ± 92.52[b] | 9 min 41 s | 260.41 ± 21.62[a] | 4 min 20 s |
| | | | SG | **1128.68 ± 173.72[a]** | 18 min 48 s | 695.4 ± 212[a] | 11 min 35 s |
| | | | IG | 760.13 ± 184.34[ab] | 12 min 40 s | 340.23 ± 44.94[a] | 5 min 40 s |
| | Blue light | | NG | 295.49 ± 71.53[a] | 4 min 55 s | 273.01 ± 55.05[a] | 4 min 33 s |
| | | | SG | 374.31 ± 106.58[a] | 6 min 14 s | 380.76 ± 95.09[a] | 6 min 20 s |
| | | | IG | 380.07 ± 69.63[a] | 6 min 20 s | 378.7 ± 63.3[a] | 6 min 18 s |
| | Red light | Optimal water regime (100% FC) | NG | 812.68 ± 106.56[ab] | 13 min 32 s | 455.1 ± 70.44[a] | 7 min 35 s |
| | | | SG | 922.24 ± 44.76[ab] | 15 min 22 s | 445.4 ± 74.36[a] | 7 min 25 s |
| | | | IG | 1005.42 ± 75.44[ab] | 16 min 45 s | 557.27 ± 137.91[a] | 9 min 17 s |
| | Blue light | | NG | 272.6 ± 57.16[a] | 4 min 32 s | 260.44 ± 48.61[a] | 4 min 20 s |
| | | | SG | 427.33 ± 25.07[a] | 7 min 7 s | 399.98 ± 24.96[a] | 6 min 39 s |

| | | | | |
|---|---|---|---|---|
| IG | 446.57 ± 50.07[a] | 7 min 26 s | 396.09 ± 41.66[a] | 6 min 36 s |

**Table S11. Time constants for stomatal conductance ($\tau_{gsw}$) and net $CO_2$ assimilation rate ($\tau_A$) during step changes in light intensity and quality in pearl millet.** We estimated in rice non-grafts (NG), self-grafts (SG) and intra-species grafts (IG), under saturated water regime of 160 % of soil water field capacity (FC) and at optimal water regime of 100 % FC at 38 and 43 days after grafting, the time constants (the time required to reach 63 % of the steady-state maximum) for stomatal conductance ($g_{sw}$) and net $CO_2$ assimilation ($A$) following step changes in light intensity (red light; from 100 to 1500 μmol $m^{-2}$ $s^{-1}$) and a change in light quality (blue light; from 1500 μmol $m^{-2}$ $s^{-1}$ red light to 1500 μmol $m^{-2}$ $s^{-1}$ composed of 90 % red and 10 % blue light). Data are means ± SEM ($n$ = 3 - 6). Data were analyzed using two-way ANOVA linear model with graft type, water regime, and their interactions as the main explanatory factors. Difference between the groups were detected by Tukeys HSD test for multiple comparisons at $p \leq 0.05$. Different lowercase superscript letters within columns indicate significant differences ($p \leq 0.05$) between graft as determined by Tukey's HSD post-hoc test. DAG, stands for days after grafting.

| Timepoint | Light | Regime | Graft | $\tau_{gsw}$ (S) | Duration (min) | $\tau_A$ (S) | Duration (min) |
|---|---|---|---|---|---|---|---|
| 38 DAG | Red light | Saturated water regime (160 % FC) | NG | $581.62 \pm 38.97^{a}$ | 9 min 41 s | $334.71 \pm 37.34^{ab}$ | 5 min 34 s |
| | | | SG | $854.19 \pm 80.38^{a}$ | 14 min 14 s | $580.39 \pm 103.39^{a}$ | 9 min 40 s |
| | | | IG | $636.4 \pm 49.87^{a}$ | 10 min 36 s | $271 \pm 23.86^{b}$ | 4 min 30 s |
| | Blue light | | NG | $226.55 \pm 24.06^{a}$ | 3 min 46 s | $387.92 \pm 12.71^{ab}$ | 6 min 27 s |
| | | | SG | $287.31 \pm 47.19^{a}$ | 4 min 47 s | $384.57 \pm 14.39^{ab}$ | 6 min 24 s |
| | | | IG | $217.33 \pm 38.44^{a}$ | 3 min 37 s | $268.74 \pm 48.98^{b}$ | 4 min 28 s |
| | Red light | Optimal water regime (100% FC) | NG | $577.83 \pm 87.32^{a}$ | 9 min 37 s | $350.96 \pm 59.18^{ab}$ | 5 min 50 s |
| | | | SG | $646.32 \pm 70.93^{a}$ | 10 min 46 s | $404.65 \pm 71.09^{ab}$ | 6 min 44 s |
| | | | IG | $753.21 \pm 75.45^{a}$ | 12 min 33 s | $302.05 \pm 7.08^{ab}$ | 5 min 2 s |
| | Blue light | | NG | $238.64 \pm 23.08^{a}$ | 3 min 58 s | $405.5 \pm 34.34^{a}$ | 6 min 45 s |
| | | | SG | $264.6 \pm 18.04^{a}$ | 4 min 24 s | $358.6 \pm 19.88^{ab}$ | 5 min 58 s |
| | | | IG | $278.66 \pm 37.37^{a}$ | 4 min 38 s | $267.12 \pm 43.25^{b}$ | 4 min 27 s |
| 43 DAG | Red light | Saturated water regime (160 % FC) | NG | $528.09 \pm 30.89^{a}$ | 8 min 48 s | $387.7 \pm 31.47^{a}$ | 6 min 27 s |
| | | | SG | $644.92 \pm 80.28^{a}$ | 10 min 44 s | $389.76 \pm 31.48^{a}$ | 6 min 29 s |
| | | | IG | $689.61 \pm 63.53^{a}$ | 11 min 29 s | $348.79 \pm 76.09^{a}$ | 5 min 48 s |
| | Blue light | | NG | $222.01 \pm 27.65^{b}$ | 3 min 42 s | $316.29 \pm 24.78^{a}$ | 5 min 16 s |
| | | | SG | $362.57 \pm 38.64^{a}$ | 6 min 2 s | $410.32 \pm 40.33^{a}$ | 6 min 50 s |
| | | | IG | $258.58 \pm 40.1^{ab}$ | 4 min 18 s | $408.21 \pm 36.99^{a}$ | 6 min 48 s |
| | Red light | Optimal water regime (100% FC) | NG | $529.84 \pm 46.13^{a}$ | 8 min 49 s | $358.61 \pm 51.05^{a}$ | 5 min 58 s |
| | | | SG | $543.32 \pm 71.8^{a}$ | 9 min 3 s | $410.63 \pm 38.86^{a}$ | 6 min 50 s |
| | | | IG | $440.71 \pm 134.99^{a}$ | 7 min 20 s | $267.83 \pm 59.5^{a}$ | 4 min 27 s |
| | Blue light | | NG | $183.76 \pm 15.53^{b}$ | 3 min 3 s | $289.42 \pm 40.35^{a}$ | 4 min 49 s |
| | | | SG | $236.79 \pm 36.16^{ab}$ | 3 min 56 s | $343.35 \pm 35.77^{a}$ | 5 min 43 s |

| | | | | | |
|---|---|---|---|---|---|
| | IG | 167.43 ± 23.92[b] | 2 min 47 s | 356.44 ± 77.32[a] | 5 min 56 s |

**Table S12. Two-way ANOVA results on time constants for stomatal conductance ($\tau_{gsw}$) and net $CO_2$ assimilation rate ($\tau_A$) during step changes in light intensity and quality in rice and pearl millet.** We estimated in rice and pearl millet non-grafts (NG), self-grafts (SG) and intra-species grafts (IG), under saturated water regime of 160 % of soil water field capacity (FC) and at optimal water regime of 100 % FC at 38 and 43 days after grafting, the time constants (the time required to reach 63 % of the steady-state maximum) for stomatal conductance ($g_{sw}$) and net $CO_2$ assimilation ($A$) following step changes in light intensity (red light; from 100 to 1500 µmol $m^{-2}$ $s^{-1}$) and a change in light quality (blue light; from 1500 µmol $m^{-2}$ $s^{-1}$ red light to 1500 µmol $m^{-2}$ $s^{-1}$ composed of 90 % red and 10 % blue light). Data are means ± SEM ($n$ = 3 - 6). Data were analyzed using two-way ANOVA linear model with graft type, water regime, and their interactions as the main explanatory factors. Statistically significant effects are indicated in bold. DAG, stands for days after grafting; Df(n,d), stands for the degree of freedom, numerator, and denominator; F stands for F test value; P stands for probability value in at ≤ 0.05.

| **Plant** | Light | Timepoint | Source of variation | $\tau_A$ | | | $\tau_{gsw}$ | | |
|---|---|---|---|---|---|---|---|---|---|
| | | | | Df(n,d) | F | P | Df(n,d) | F | P |
| Rice | Red light | 38 DAG | Graft | 2,20 | 1.824 | 0.187 | 2,20 | 1.505 | 0.246 |
| | | | Regime | 1,20 | 0.383 | 0.543 | 1,20 | 0.310 | 0.584 |
| | | | Graft*Regime | 2,20 | 0.773 | 0.475 | 2,20 | 0.428 | 0.658 |
| | Blue light | | Graft | 2,20 | 0.422 | 0.662 | 2,20 | 1.340 | 0.284 |
| | | | Regime | 1,20 | 2.695 | 0.116 | 1,20 | 1.363 | 0.257 |
| | | | Graft*Regime | 2,20 | 1.010 | 0.382 | 2,20 | 2.494 | 0.108 |
| | Red light | 43 DAG | Graft | 2,16 | 2.423 | 0.120 | 2,16 | 2.770 | 0.093 |
| | | | Regime | 1,16 | 0.434 | 0.519 | 1,16 | 1.296 | 0.272 |
| | | | Graft*Regime | 2,16 | 3.239 | 0.066 | 2,16 | 1.166 | 0.337 |
| | Blue light | | Graft | 2,16 | 2.578 | 0.107 | 2,16 | 2.085 | 0.157 |
| | | | Regime | 1,16 | 0.043 | 0.838 | 1,16 | 0.149 | 0.704 |
| | | | Graft*Regime | 2,16 | 0.126 | 0.882 | 2,16 | 0.197 | 0.823 |
| Pearl millet | Red light | 38 DAG | Graft | 2,22 | 4.704 | **0.020** | 2,22 | 2.140 | 0.141 |
| | | | Regime | 1,22 | 0.510 | 0.482 | 1,22 | 0.132 | 0.720 |
| | | | Graft*Regime | 2,22 | 1.497 | 0.246 | 2,22 | 1.551 | 0.234 |
| | Blue light | | Graft | 2,22 | 10.142 | **0.000** | 2,22 | 0.883 | 0.428 |
| | | | Regime | 2,22 | 0.058 | 0.812 | 1,22 | 0.116 | 0.737 |
| | | | Graft*Regime | 1,22 | 0.473 | 0.629 | 2,22 | 0.762 | 0.479 |
| | Red light | 43 DAG | Graft | 2,18 | 1.203 | 0.323 | 2,18 | 0.396 | 0.678 |
| | | | Regime | 1,18 | 1.062 | 0.316 | 1,18 | 2.842 | 0.109 |
| | | | Graft*Regime | 2,18 | 0.488 | 0.622 | 2,18 | 1.461 | 0.258 |
| | Blue light | | Graft | 2,18 | 1.732 | 0.205 | 2,18 | 6.058 | **0.009** |

| | | | | | | | |
|---|---|---|---|---|---|---|---|
| | Regime | 1,18 | 2.263 | 0.150 | 1,18 | 7.139 | **0.016** |
| | Graft*Regime | 2,18 | 0.007 | 0.993 | 2,18 | 1.370 | 0.279 |

**Table S13. Two-way ANOVA results on the photosynthetic-stomatal coordination index ($\tau_{gsw}/\tau_A$) during step changes in light intensity and quality in rice and pearl millet**. We calculated the coordination index as the ratio between the time constants for stomatal conductance ($\tau_{gsw}$) and net $CO_2$ assimilation rate ($\tau_A$) in rice and pearl millet's non-grafts (NG), self-grafts (SG) and intra-species grafts (IG) under a saturated water regime of 160 % of soil water field capacity (FC) and at optimal water regime of 100 % FC at 38 and 43 days after grafting. Time constants represent the time required to reach 63 % of the steady-state maximum following step changes in light intensity (red light; from 100 to 1500 µmol $m^{-2}$ $s^{-1}$) and a change in light quality (blue light; from 1500 µmol $m^{-2}$ $s^{-1}$ red light to 1500 µmol $m^{-2}$ $s^{-1}$ composed of 90 % red and 10 % blue light). Data are means ± SEM (*n* = 3 - 6). Data were analyzed using two-way ANOVA linear model with graft type, water regime, and their interactions as the main explanatory factors. Statistically significant effects are indicated in bold. DAG, stands for days after grafting; Df(n,d), stands for the degree of freedom, numerator, and denominator; F stands for F test value; P stands for probability value in at ≤ 0.05.

| Plant | Timepoint | Source of variation | Coordination Index | | | | | |
|---|---|---|---|---|---|---|---|---|
| | | | Red light | | | Blue light | | |
| | | | Df(n,d) | F | P | Df(n,d) | F | P |
| Rice | 38 DAG | Graft | 2,20 | 0.511 | 0.608 | 2,20 | 1.079 | 0.359 |
| | | Regime | 1,20 | 3.622 | 0.072 | 1,20 | 1.810 | 0.193 |
| | | Graft*Regime | 2,20 | 0.016 | 0.984 | 2,20 | 1.533 | 0.240 |
| | 43 DAG | Graft | 2,16 | 0.486 | 0.624 | 2,16 | 1.319 | 0.295 |
| | | Regime | 1,16 | 0.072 | 0.792 | 1,16 | 1.027 | 0.326 |
| | | Graft*Regime | 2,16 | 1.963 | 0.173 | 2,16 | 0.200 | 0.820 |
| Pearl millet | 38 DAG | Graft | 2,22 | 7.941 | **0.003** | 2,22 | 6.198 | **0.007** |
| | | Regime | 1,22 | 0.096 | 0.759 | 1,22 | 0.541 | 0.470 |
| | | Graft*Regime | 2,22 | 0.152 | 0.860 | 2,22 | 0.446 | 0.646 |
| | 43 DAG | Graft | 2,18 | 1.800 | 0.194 | 2,18 | 2.033 | 0.160 |
| | | Regime | 1,18 | 0.795 | 0.384 | 1,18 | 1.132 | 0.301 |
| | | Graft*Regime | 2,18 | 1.423 | 0.267 | 2,18 | 1.297 | 0.298 |
